# Predicted Future Changes in the Mean Seasonal Carbon Cycle Due to Climate Change

**DOI:** 10.1101/2024.05.20.594964

**Authors:** Mauro Morichetti, Elia Vangi, Alessio Collalti

**Affiliations:** National Research Council of Italy, Forest Modelling Lab., Institute for Agriculture and Forestry Systems in the Mediterranean (CNR-ISAFOM), Via Madonna Alta 128, 06128 Perugia, Italy; geoLAB - Laboratory of Forest Geomatics, Dept. of Agriculture, Food, Environment and Forestry, Università degli Studi di Firenze, Via San Bonaventura 13, 50145 Firenze; National Biodiversity Future Center (NBFC), 90133 Palermo, Italy

**Keywords:** Carbon Cycle, Climate Change, Process-Based Model, Mean Seasonal Cycle, Forest Ecosystems

## Abstract

Through photosynthesis, forests absorb annually large amounts of atmospheric CO_2_. However, they also release CO_2_ back through respiration. These two, opposite in sign, large fluxes determine, much of the carbon that is stored or released back to the atmosphere. The mean seasonal cycle (MSC) is an interesting metric that associates phenology and carbon (C) partitioning-allocation analysis within forest stands. Here we applied the 3D-CMCC-FEM model and analyzed its capability to represent the main C-fluxes, by validating the model against observed data, questioning if the sink/source mean seasonality is influenced under two scenarios of climate change, in five contrasting European forest sites. We found the model has, under current climate conditions, robust predictive abilities in estimating NEE. Model results also predict a consistent reduction of the forest’s capabilities to act as a C-sink under climate change and stand-ageing at all sites. Such a reduction is predicted despite the number of annual days of C-sink in evergreen forests increasing over the years, indicating a consistent downward trend. Similarly, deciduous forests, despite maintaining a relatively stable number of C-sink days throughout the year and over the century, show a reduction in their overall annual C-sink capacity. Overall, both types of forests at all sites show a consistent reduction in their future mitigating potential.

## 1. Introduction

Forests play a pivotal role in the biosphere-atmosphere feedback by annually absorbing large amounts of atmospheric CO_2_ through photosynthesis (GPP; ∼150 PgC yr^─1^) and releasing it back because of e.g., ecosystem respiration (R_eco_), a relatively close amount yet not necessarily equal, that varies year by year [1]–[3]. The net ecosystem exchange (NEE) of CO_2_ between ecosystems and the atmosphere is the net balance between these two gross fluxes opposite in signs, and it governs much of the overall terrestrial annual net carbon (C) budget. Imbalances between CO_2_ sources (even including carbon lost by fires and other processes) and sinks directly increase or decrease atmospheric CO_2_ levels [4]. Terrestrial ecosystems ─ and forests in particular ─ are contributing substantially to climate change mitigation, provided that they are C-sinks and not C-sources [5]. Forests that might absorb more than they emit are commonly considered carbon sinks (with NEE negative in sign), while if they emit more than they absorb are considered as carbon sources (with NEE positive in sign). The magnitude of this exchange of CO_2_, however, is subject to substantial variability and trends, in large part as a response to variations and trends in climate [6]. Indeed, forests have been shown to be extremely sensitive to changes in environmental conditions (e.g., climate, seasonality, atmospheric CO_2_ concentration, nitrogen deposition), to ageing [7], [8] and to disturbances [9], including management practices [10], [11] which can control both photosynthesis and respiration. Therefore, estimating NEE, GPP, and R_eco_ is a key step for better understanding the underlying mechanisms constraining ecosystem functioning [12].

Europe and the Mediterranean are expected to become in the near future ‘Hot Spot’ of climate change [13]–[16]. Literature reports that under climate change scenarios forests are expected to grow faster, to mature earlier but also to die younger, curtailing their life-span [17], because of, mainly, warming and increased atmospheric CO_2_ concentration (the so-called ‘CO_2_-fertilization effect’) [18]. Conversely, there is a general lack of evidence and knowledge on how, overall, forest ecosystems will, on the whole, react to climate change. Forest carbon balance will be impacted by climate change because different main processes are impacted, which, in turn, may react and respond differently to climate change also because they are vulnerable and sensitive to separate environmental drivers. Indeed, while an extensive line of evidence shows as the increased availability of CO_2_ may amplify the photosynthetic rate and assimilation capacity [19] such an increase is largely debated since there is no evidence that such positive changes will generally continue indefinitely [20], [21]. Similarly, the effects of warming are largely discussed because while it is documented that some species may take advantage of a longer vegetation season (e.g., deciduous species), there are also negative effects linked to warming as heat waves and the often associated drought events [22] [23], including late frost [24] and disturbances [25], can be detrimental to growth till tree survival. In addition, there is a general concern that the changing temperature response of respiration turns boreal forests from carbon sinks into carbon sources [26]. Indeed, warming has also been found to accelerate both autotrophic as well as heterotrophic respiration (the two components of ecosystem respiration), meaning that increased temperature may lead forests to release more carbon, potentially more than absorbed annually [27]. Conversely, drought has been shown to reduce microbial respiration and then heterotrophic respiration [28]. How these processes (i.e., photosynthesis and ecosystem respiration) will be impacted by climate change annually will determine much of the future forest annual C-budget. The Mean Seasonal Cycle (MSC) metric, which reflects the average distribution of flux (i.e., NEE, GPP and R_eco_) throughout the days of a year, is an insightful measure linking phenology with carbon partitioning and allocation within seasonal climatic variability. By capturing the typical fluctuations in a specific region due to changing seasons, the MSC highlights the expected seasonal changes in climate data, averaged over many years to smooth out anomalies and emphasize the regular, cyclical nature of these changes. Many studies [29]–[32] have indeed shown that climate change will impact both the phenology by changing the date for the beginning and the end of the growing season as well as by changing the shape of the Leaf Area Index (LAI) distribution over the year which, at the same time, will influence the way, among the other things, when photosynthesis can start and how recent and old photosynthates are partitioned and used to build new tissues and to replenish the reserves used for the metabolism, as well as carbon allocation [33] and C-dynamic.

Process-based models are valuable tools to understand how and to which extent future climate change will impact these two fluxes (GPP and R_eco_) in the MSC, being both processes controlled by warming and changes in precipitation regime and atmospheric CO_2_ concentration [34], [35]. Here, we at first applied and validated under current observed climate conditions the ‘*Three Dimensional - Coupled Model Carbon Cycle - Forest Ecosystem Module*’ (3D-CMCC-FEM), a biogeochemical, biophysical process-based forest ecosystem model designed to simulate carbon, nitrogen, and water cycle in forest ecosystems and, secondly, under climate change conditions. Specifically, we question and analyze: 1) the capability of the 3D-CMCC-FEM to represent under the current climate the main C-fluxes governing C-cycle in terms of net ecosystem exchange (NEE), gross primary production (GPP) and ecosystem respiration (R_eco_), by validating the model against independent data from the Fluxnet network; and, provided that the model is close to the observed data; 2) how, and if, the sink/source mean seasonality will be influenced under two locally bias-corrected scenarios of warming (RCP 2.6 and 6.0) and atmospheric CO_2_ enrichment from three CMIP5 Earth System Models, within ISIMIP-PROFOUND initiative, in five well-studied and long-monitored contrasting forest sites (three evergreens and two deciduous) on a longitudinal transect through Europe up to the end of the century.

## 2. Materials and Method

### 2.1. Model description (3D-CMCC-FEM ‘v.5.6’)

The *’Three Dimensional - Couple Model Carbon Cycle - Forest Ecosystem Module*’ (hereafter ‘3D-CMCC-FEM’) is a biogeochemical, biophysical, process-based forest ecosystem model (see [10], [11], [36]–[44] and reference therein). The model is designed to simulate carbon, nitrogen and water cycles in forest ecosystems at commonly 1-hectare spatial resolution and the main eco-physiological processes (e.g., photosynthesis) at daily temporal resolution. The most recent code versions since Collalti et al. [17] adopt the biogeochemical photosynthesis model of Farquhar, von Caemmerer and Berry [45] to compute gross primary productivity (GPP). The biogeochemical photosynthesis model is parameterized as in Bernacchi et al. [46], [47] and temperature acclimation for leaves as in Kattge and Knorr [48]. The 3D-CMCC-FEM considers, as in De Pury and Farquhar [49], light interception, reflection, transmission, and assimilation (and leaf respiration) for both sun and shaded leaves. Autotrophic respiration (R_A_) is computed mechanistically following the ‘*Growth and Maintenance Respiration Paradigm*’ (GMRP; [50]), which is divided into the metabolic costs for synthesizing new tissues (growth respiration, R_G_) and the metabolic costs for maintaining the existing ones (maintenance respiration, R_M_). In 3-D-CMCC-FEM, the maintenance respiration is based on Nitrogen amount (a fixed fraction of carbon mass varying between the six tree compartments) and is temperature-controlled by a standard Arrhenius relationship [36]. ‘Type I’ and ‘Type II’ acclimation of respiration to temperature (i.e., short- and long-term acclimation; [10], [39], [51]) are also accounted for. Any imbalance between carbon assimilation and carbon losses because of plants’ respiration is buffered by a seventh pool, the Non-Structural Carbon pool (NSC; starch and sugars undistinguished), that has priority in the carbon allocation all over the year. The net primary production (NPP) is the GPP minus R_A_. Biomass production (BP) is the amount of NPP not used for replenishing the NSC pool. Indeed, other forms of non-structural carbon losses (e.g., biogenic volatile organic compounds, BVOCs, or root exudates to mycorrhizas) are not accounted for by the model. The phenological scheme, as well as the carbon partitioning-allocation scheme, distinguished for deciduous and evergreen tree species, is in-depth described in Collalti et al. [10], [36], [39] and [52]. Heterotrophic respiration follows a BIOME-BGC-like approach (which follows the CENTURY model; [53], [54]) distinguishing decomposition for litter and soil pools with, each, four different conceptual pools characterized by different decomposability degrees (i.e., fast, medium, slow and a recalcitrant carbon pool) [39], [55]. Altogether, litter and soil decomposition emissions form heterotrophic respiration (R_H_), which, summed up to R_A_, constitutes ecosystem respiration (R_eco_). Net ecosystem exchange (NEE) is equal to R_eco_ minus GPP. Therefore, negative values indicate carbon uptake from the atmosphere (i.e., the system acts as a C-sink, NEE < 0), and positive values indicate carbon release (i.e., the system acts as a C-source, NEE > 0). The 3D-CMCC-FEM’s sensitivity to its 54 species-specific parameters and how it varies along the forest development and under climate change is described in Collalti et al. [17].

In the present study, we used version 5.6 [41], which slightly differs from v.5.5, described in Collalti et al. [8] and Dalmonech et al. [11], for a new scheme (and relative parameterization) for sapwood and live wood turnover and dynamics and some additional new forest management schemes (not used here) described in Testolin et al. [43].

### 2.2. Case study areas

Five case studies have been selected as representative of the main European forest species (and climate) and at the same time because of long-monitored sites and part of the Fluxnet network [56], the ISIMIP (Inter-Sectoral Impact Model Intercomparison Project, [57]; https://www.isimip.org/) initiative and the PROFOUND database [42], [58], [59]: the temperate European beech (*Fagus sylvatica* L.) forest of Collelongo, Italy (IT-Col), and of Sorø, Denmark (DK-Sor), the maritime pine (*Pinus pinaster* Ait.) forest of Le Bray, France (FR-Lbr), the boreal Scots pine (*Pinus sylvestris* L.) forest of Hyytiälä, Finland (FI-Hyy), and the temperate Norway spruce (*Picea abies* (L.) H. Karst) forest of Bílý Křìž, Czech Republic (CZ-Bk1) (Figure 1). Stand characteristics are described in Table 1.

**Figure 1.**
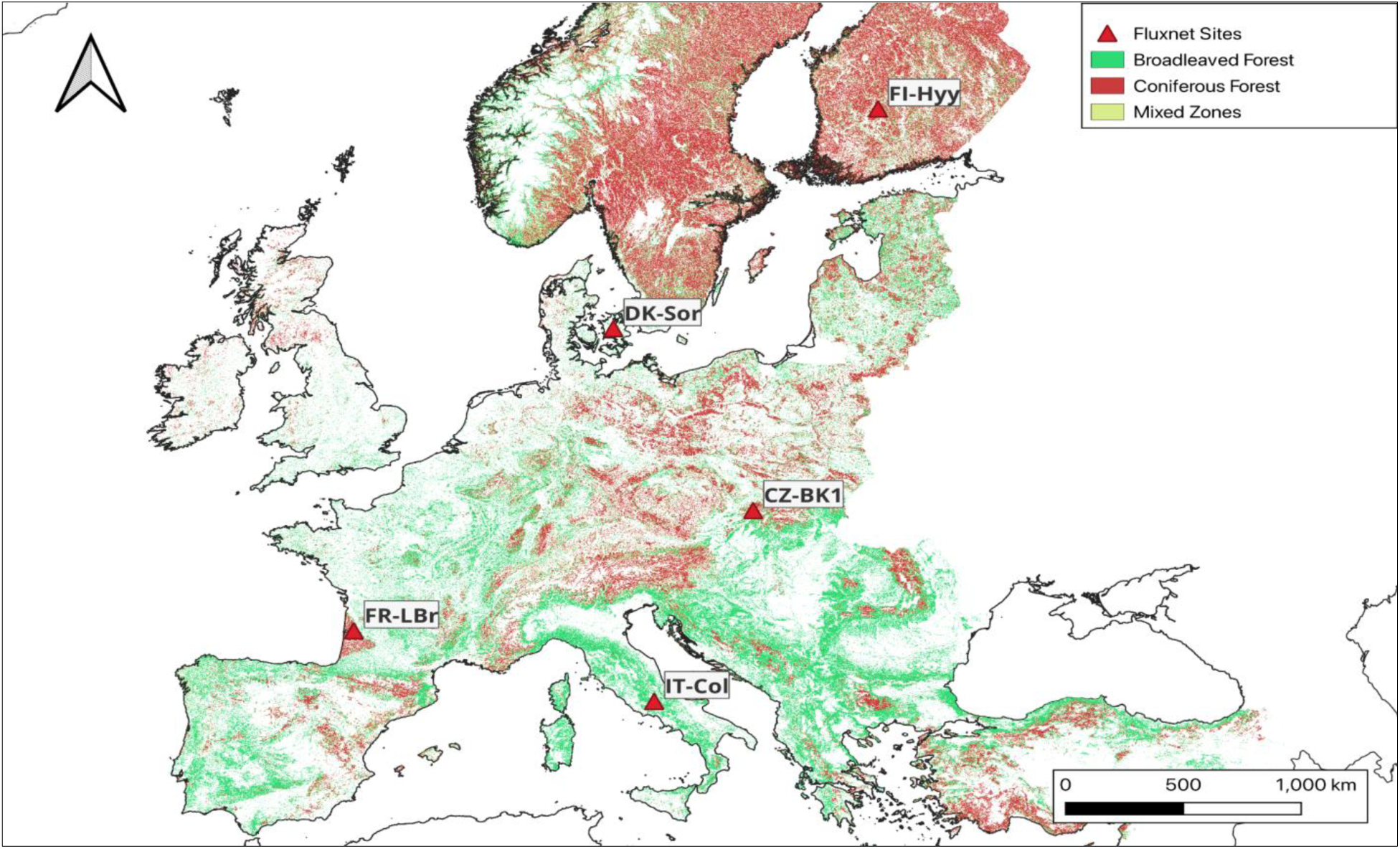
Forest classification for broadleaved and coniferous forests at the European level in the spatial resolution of 100 m (ESA, 2020). Location of Fluxnet sites considered to assess the 3D-CMCC-FEM.

**Table 1.** Site description and stand initialization data are used in simulations with data from the PROFOUND database (Reyer et al., 2020). Model initialization data (i.e., diameter at breast height - DBH, tree height, stand age, and stand density) correspond to the stand conditions of the first year in the historical simulations.

| Fluxnet<br>Code | Nation | Site Name | Coordinate |  |  | Climate | Species<br>Composition | Period of simulation<br>and climate data |  | Mean<br>DBH<br>(cm) | Tree<br>Height<br>(m) | Mean<br>Stand<br>Age<br>(years) | Stand<br>Density<br>(n. trees<br>ha<sup>-1</sup>) |
| --- | --- | --- | --- | --- | --- | --- | --- | --- | --- | --- | --- | --- | --- |
|  |  |  | Lat<br>(°) | Lon<br>(°) | Elevation<br>(m) |  |  | Historical<br>(observed) | Historical +<br>RCP<br>(modelled) |  |  |  |  |
| IT-Col | Italy | Collelongo | 41.8 | 13.6 | 1560 | Temperate | *Fagus*<br>*sylvatica* | 1997-2014 | 1997-2099 | 20.2 | 17.3 | 105 | 900 |
| FR-LBr | France | Le Bray | 44.7 | -0.8 | 61 | Temperate | *Pinus*<br>*pinaster* | 1996 -<br>2008 | 1997-2099 | 26.7 | 17.8 | 26 | 614 |
| FI-Hyy | Finland | Hyytiälä | 61.9 | 24.3 | 181 | Boreal | *Pinus*<br>*sylvestris* | 1996-2014 | 1997-2099 | 13 | 11.3 | 35 | 870 |
| CZ-Bk1 | Czech<br>Republic | Bílý Kříž | 49.3 | 18.3 | 875 | Temperate | *Picea abies* | 2000-2008 | 1997-2099 | 9.1 | 7.5 | 19 | 2388 |
| DK-Sor | Denmark | Sorø | 55.3 | 11.4 | 40 | Boreal | *Pinus*<br>*sylvestris* | 1996 -<br>2008 | 1997-2099 | 24.3 | 21 | 75 | 353 |

### 2.3. Input, meteorological data and climate change scenarios

To run, 3D-CMCC-FEM needs a set of input data from state variables representing the stand at the beginning of simulations and that account for structural characteristics (e.g., tree height, average age, diameter at breast height; see Table 1) as well as carbon and nitrogen pools (e.g., stem carbon and nitrogen); meteorological forcing data (e.g., daily maximum and minimum temperature, daily precipitation); annual atmospheric CO_2_ concentrations; and species-specific parameters (e.g., maximum stomatal conductance). All the data used to initialize the model in the present study for the five stands come from the ISIMIP initiative and the PROFOUND database [58]. More specifically, daily observed meteorological data for model validation come from the Fluxnet2015 Dataset [56], while daily modeled historical (1997-2005) and future climate scenarios (2006-2099) are those from the ‘ISIMIP 2bLBC’ experiments (‘2b experiments Locally Bias Corrected’) coming from three different Earth System Models (ESMs; GFDL-ESM2M, IPSL-CM5A-LR, and MIROC5, respectively) based on the Climate Model Intercomparison Project 5 (CMIP5) driven by two Representative Concentration Pathways (RCPs) of atmospheric greenhouse gas concentration trajectories, namely RCP 2.6 and RCP 6.0. [60], [61]. The ISIMIP 2bLBC have the same structure as those in the 2b experiments but have been corrected by improving the method described in [62] and subsequently by the methods described in Frieler et al. [57], and Lange [63] using the observed meteorology at the local level [58]. Therefore, the 2bLBC climate data represent the more consistent and closer modeled climate data with the observational data. The annual atmospheric CO_2_ concentrations for the historical period are based on Meinshausen et al. [64] and have been extended for the period from 2006 to 2015 with data from Dlugokencky and Tans [65]. Values specific for each RCP for the period 2016 to 2099 are also based on Meinshausen et al. [64] and were used within the Farquhar, von Caemmerer and Berry [45] photosynthesis model with values varying, at the end of this century, from 421.4 μmol mol^—1^ (RCP 2.6) to 666.4 μmol mol^—1^ (RCP 6.0), respectively. NEE, GPP, and R_eco_ data, for model validation come from the Fluxnet2015 Dataset [56]. Other variables have been validated at these forest stands in the past (although using slightly different model versions) and described in Collalti et al. [8], [10], [37], Marconi et al. [41], Mahnken et al. [43], Dalmonech et al. [11].

### 2.4. Model runs, validation and Mean Seasonal Cycle under climate change

The model simulations for model validation under measured forcing climate ran for CZ-Bk1 from 2000 to 2008, for IT-Col from 1997 to 2014, for FI-Hyy from 1996 to 2014, for FR-Lbr from 1996 to 2008, and for DK-Sor from 1996 to 2008 (see Table 1). For all sites model simulations under climate change scenarios began in 1997 and finished in December 2099. Model validation was performed by comparing modeled NEE, GPP, and R_eco_ against measured eddy covariance estimates (for GPP and R_eco_ using the night-time method with constant USTAR [66], as reported in the Fluxnet2015 Dataset [56].

To analyze 3D-CMCC-FEM’s capabilities to simulate NEE, GPP, and R_eco_ for daily and monthly time series, a set of commonly used statistical metrics have been applied to compare measured vs. modeled data (under observed climate forcing). To avoid considering bad quality data, a filtering procedure for quality-check has been applied; days with less than 60% of valid data were not considered and excluded both in the model and in the observed datasets. Therefore, daily NEE, GPP, and R_eco_ eddy covariance data with low-quality values (i.e., less than 0.6; [67]) were removed. Consequently, the corresponding daily modeled data were removed too. The monthly NEE, GPP, and R_eco_ values (both from eddy covariance and the model) have been aggregated from the daily ones. The common statistic we applied includes Pearson’s correlation coefficient (r), Relative Mean Bias (RMB), Normalized Root Mean Square Error (NRMSE), and Modeling Efficiency (ME)

In climate change projections, we considered the potential modifications in the ability of forest stands to absorb or emit carbon throughout the season and across the years under two different locally bias-corrected climate change scenarios, each coming from three Earth System Models. This involves averaging the daily values of the MSC of the three fluxes considered. It is noteworthy that NEE represents the equilibrium between carbon absorption by vegetation during photosynthesis and carbon release through vegetation and microbial respiration. Not only the length of the growing season but also the balance between the yearly amount of photosynthesis and R_eco_ has been shown to control much of the variability across the sites and the decades analyzed [47], [49]. This calculation is derived from the variance between GPP and R_eco_ encompassing both autotrophic respiration (R_A_, including ground components) and heterotrophic respiration (R_H_) [68].

Under climate change scenarios, we account for potential changes in the sink/source capacity of the stands during the season by averaging NEE, GPP, and R_eco_ values every ten years up to 2100 [69] and accounting for changes in the sink/source and source/sink length during the year, computed as the number of total days of the year (DoY) where a forest stand behaves as C-sink (GPP > R_eco_ with NEE < 0) or C-source (GPP < R_eco_ with NEE > 0) as described by the NEE. In addition, we also account for the changes in the DoY where NEE changes its sign at least for ten consecutive days to avoid artifact effects of pulsing, e.g., the ‘Birch effect’ on R_eco_ [70], and to account for unstable conditions and no clear source/sink and sink/source seasonal transition during the year. Therefore, we discuss changes in the MSC under the RCP 2.6 and RCP 6.0 for NEE, being the net result of opposite fluxes (i.e., GPP and R_eco_), through its changes in negative values, i.e., days in the year where NEE < 0 and describing CO_2_ uptake from the atmosphere, and positive, i.e., days in the year where NEE > 0 and describing CO_2_ release to the atmosphere. In this way, we account for potential changes that may lead to anticipations or delays in the switch from source/sink and sink/source capacity, which often happens during spring and autumn during the year. The analyses under climate change scenarios (2006 - 2099) also include the changes in the shape of the Mean Seasonal Cycle (MSC) for NEE, GPP, and R_eco_ values and for the changes (both in the absolute and the percentage values) in the annual value. Changes in MSC have been estimated on the ten-year average values using the 1998-2008 decade as a benchmarking reference. Furthermore, we account for changes in the annual values of NEE, GPP, and R_eco_ due to climate change.

## 3. Results

### 3.1. Modelled vs. observed data

The NEE, GPP, and R_eco_, as modeled by 3D-CMCC-FEM, exhibit strong correlations with the observed daily and monthly eddy covariance data and for the overall MSC across all five sites (Figure 2 and Figure A1 and Table A1, A2 and A3 in Appendix). Some slight differences are observed for the daily values for temperate European beech forests of IT-Col and DK-Sor when representing the NEE and R_eco_ between the 100 and 200 DoY (Figure 2 1.a-1.c and Figure 2 5.a-5.c). Modeled overestimations for GPP of about 5 gC m^—2^ day^—1^ at the peak of production (∼180 DoY) for both the European beech forest at IT-Col (Figure 2 - 1.b) and the maritime pine forest at FR-Lbr (Figure 2 - 2.b) was observed. The highest correlation coefficients between modeled and observed data were observed for DK-Sor and IT-Col (r = 0.97 and 0.96, respectively) for the daily NEE and GPP (Table A1, Appendix), while FI-Hyy shows the best correlation for daily R_eco_ (r = 0.94) and monthly NEE (r = 0.99) (Table A2 and Table A3).

**Figure 2.**
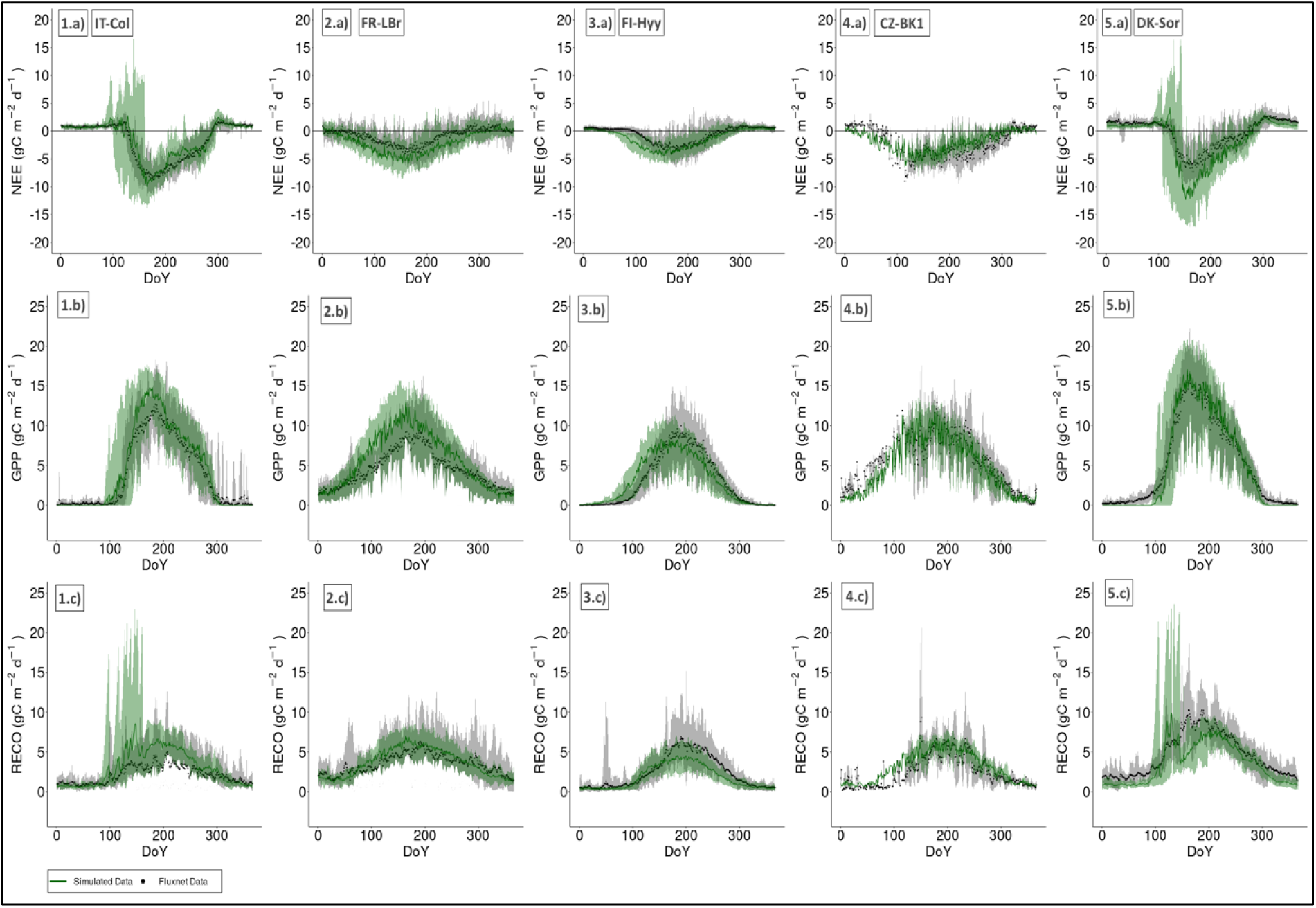
The green lines represent the modeled (a) NEE, (b) GPP and (c) R_eco_ amounts (gC m^—2^ day^—1^) per DoY (Day of Year) for the five selected case studies i.e.: 1) Collelongo – IT-Col, 2) Le Bray - FR-Lbr, 3) Hyytiälä - FI-Hyy, 4) Bílý Křìž - CZ-Bk1, and 5) Sorø - DK-Sor) compared to relative observed data (depicted as black dots) from the Fluxnet2015 Dataset (Pastorello et al., 2020). The lower and upper lines of the shaded area represent, respectively, the minimum and maximum values of the observed and modelled datasets considered.

At the IT-Col and CZ-Bk1 the model shows the best performances in simulating NEE (RMB = 0.07 and 0.14, respectively - Table A1), while the lowest values were reached by DK-Sor and IT-Col for the GPP (RMB ranging between —0.06 and 0.55, Table A2). Overall, RMB values for GPP are relatively low across all case studies and time scales. Regarding R_eco_ at FI-Hyy and DK-Sor, the model tended to underestimate both daily (RMB of -0.3 and -0.42, respectively) and monthly (-0.29 and -0.42, respectively). Conversely, at IT-Col, the model exhibited slight overestimation (RMB daily 0.82 and monthly 0.77), Table A3).

The model reports negative NRMSE values for NEE across all time scales, indicating a slight overestimation (values ranging from —0.57 to —6.59) with the exception of CZ-Bk1 site for daily NEE value (1.39), and at IT-Col site for monthly NEE value (0.99). Regarding the GPP, (Table A2) DK-Sor showed the lowest NRMSE for daily and monthly values (0.19 and 0.16, respectively). At the opposite, at CZ-Bk1 and FR-Lbr forests, the model displays the highest NRMSE for daily (1.21) and monthly (0.40) values, respectively. Last, Table A3 shows the validation results for R_eco_. At FR-LBR the highest accuracy and precision were reported with the lowest NRMSE for daily and monthly values (0.19 and 0.16 respectively). For the other sites, we found almost the same model capability described for the above fluxes (i.e., NEE and GPP) with a lower correlation and a slightly higher error in terms of daily R_eco_ at CZ-Bk1 (NRMSE = 1.16). ME exhibits values close to one across all time scales and sites for NEE, with values ranging from —0.10 (FR-LBr) to 0.92 (IT-Col) for daily NEE values and —0.15 (FR-LBr) to 0.95 (IT-Col) for monthly values. The modeled GPP has a similar trend of the observed NEE with values ranging from 0.28 for FR-LBr to 0.97 for DK-Sor for daily values and FR-LBr = 0.34 to DK-Sor = 0.92 for monthly values (Table A2). The R_eco_ achieved lower performances than GPP and NEE in terms of ME; Table A3 displays values between 1.17 for IT-Col to 0.75 for FI-Hyy and IT-Col = —0.64 to FI-Hyy = 0.77, for daily and monthly time scale, respectively.

Finally, the lowest MAE was found at IT-Col, for NEE (MAE = 0.67 gC m^—2^ day^—1^ and MAE = 11.42 gC m^—2^ month^—1^) and GPP (MAE = 0.91 gC m^—2^ day^—1^ and MAE = 20.68 gC m^—2^ month^—1^), while the lowest MAE for R_eco_ was for FR-LBr (MAE = 0.71 gC m^—2^ day^—1^ and MAE = 18.01 gC m^—2^ month^—1^) (Table A1, Table A2, and Table A3 in Appendix).

### 3.2. Mean seasonal NEE cycle under climate change scenario

Figure 3 and Figure 4 display the 10-year average NEE seasonal cycle under climate scenarios (i.e., RCP 2.6 and RCP 6.0) for the five case studies. Overall, across all the sites and scenarios considered, there is a consistent reduction in the absolute NEE over time (i.e., NEE is ‘less negative’ showing a reduction in the sink capacity) with changes in the source/sink (NEE becomes negative and the site turns to C-sink) and sink/source (NEE becomes positive and the site turns to C-source) switch over the year(s). However, RCP 2.6 generally exhibits lower reductions in annual and mean seasonal NEE when compared to RCP 6.0 across most study sites and time intervals. The loss in the modeled sink capacity is because the R_eco_ increases more than GPP does and the differences between the scenarios are related to an increase in R_eco_ higher than for GPP under RCP 6.0.

**Figure 3.**
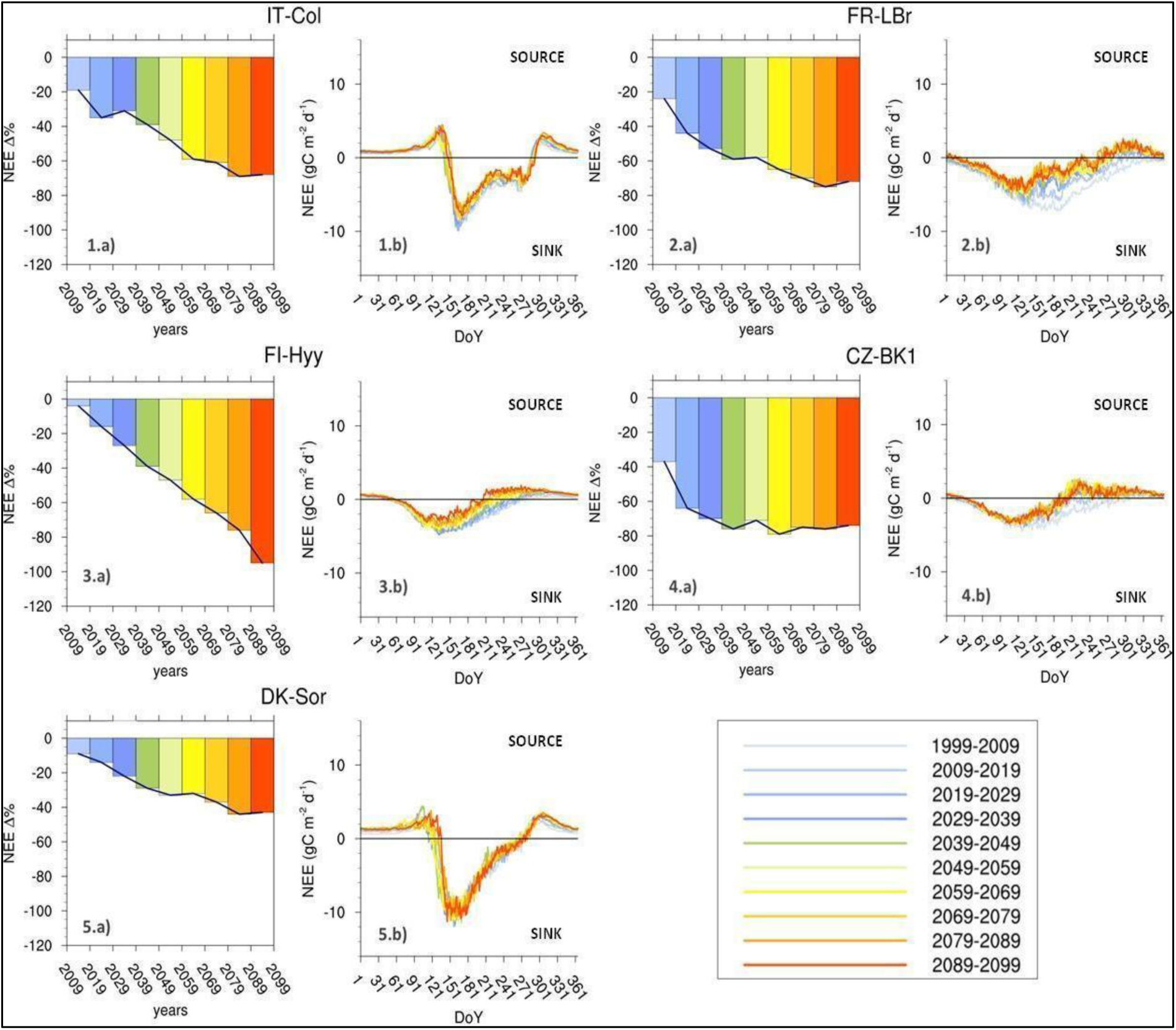
10-year average NEE seasonal cycle under the RCP 2.6 climate scenario for 5 case studies selected, i.e.: 1) Collelongo - IT-Col, 2) Le Bray - FR-LBr, 3) Hyytiälä - FI-Hyy, 4) Bílý Kříž - CZ-Bk1, and 5) Sorø - DK-Sor). The histograms (a) represent the annual NEE variation (%) from the first decade taken as a benchmark of simulation (1999-2009). The xy plots (b) show the Mean Seasonal NEE Cycle of daily values (gC m^—2^ day^—1^).

**Figure 4.**
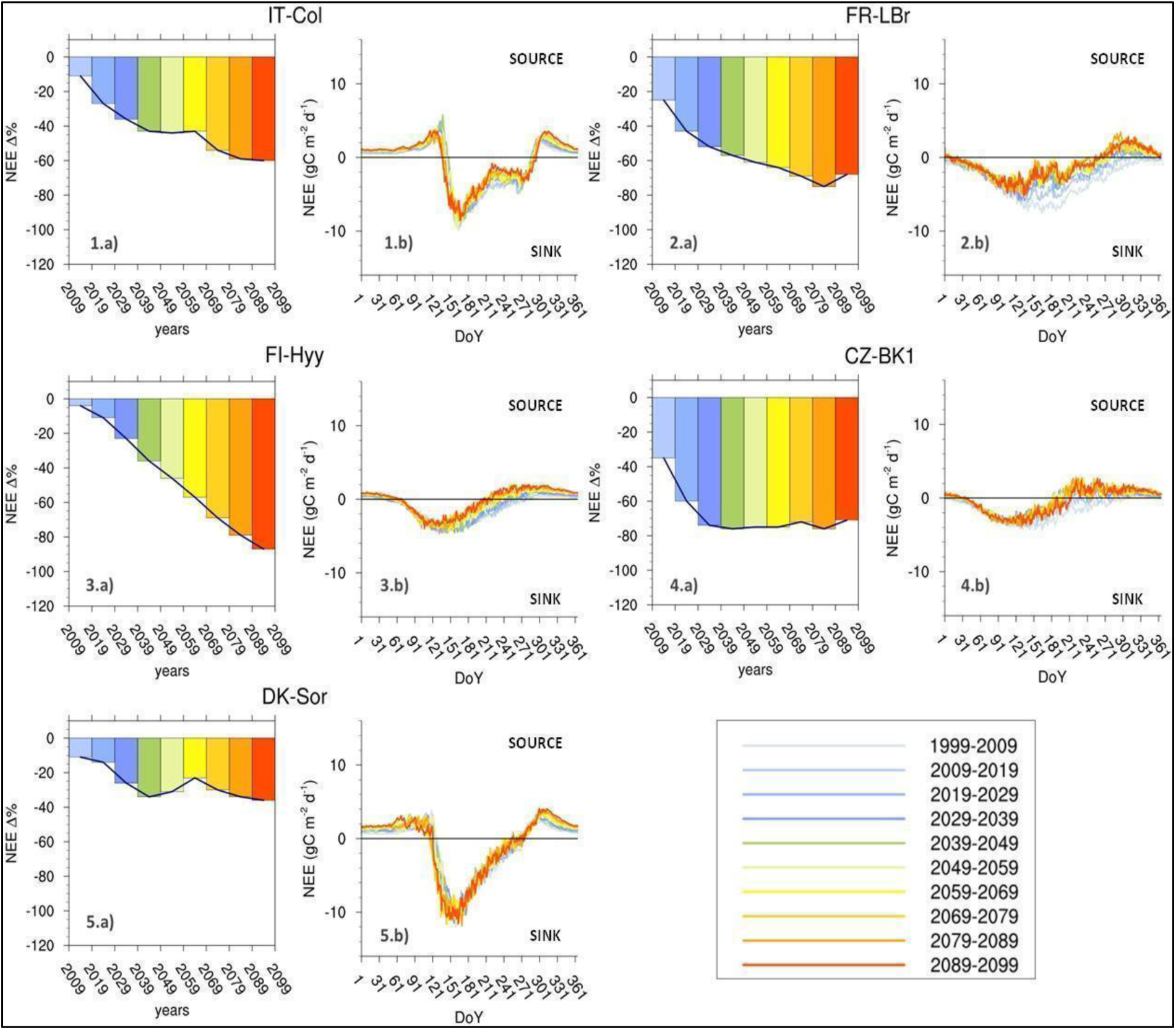
10-year average NEE seasonal cycle under the RCP 6.0 climate scenario for 5 case studies selected, i.e.: 1) Collelongo - IT-Col, 2) Le Bray - FR-LBr, 3) Hyytiälä - FI-Hyy, 4) Bílý Kříž - CZ-Bk1, and 5) Sorø - DK-Sor). The histograms (a) represent the annual NEE variation (%) from the first decade taken as a benchmark of simulation (1999-2009). The xy plots (b) show the Mean Seasonal NEE Cycle of daily values (gC m^—2^ day^—1^).

The rate of decrease in the annual NEE varies among locations and forest species. For example, the beech forests at IT-Col and DK-Sor show a reduction in the NEE (i.e., the site is less C-sink) of about 68%, and 43% (Figure 3 and Figure 4 - panel 1 and panel 5) at the end of the century, generally exhibiting a more moderate decrease compared to evergreen sites. The Scots pine forest shows, at the end of the century, a reduction in the sink capacity of 95%, standing out as the most significant decrease in NEE (Figure 3 and Figure 4 – panel 3). Over time, the rate of decline shows a tendency to speed up, hinting a reduction in the capacity of the carbon sink because, despite an overall increase in the GPP (Figure A2 and A3), the increased ecosystem respiration due to increased temperatures (Figure A4 and A5). By the end of the century, substantial reductions in NEE, i.e., the sites become less C-sink, across all locations and scenarios were simulated, with some locations experiencing over 70-90% reduction compared to the 1999-2009 decade (Table A4).

The GPP, similar to NEE, generally increases across forest types and scenarios over time, albeit with varying degrees (Fig A2, A3). Specifically, RCP 6.0 shows higher growth rates, particularly in later years (2059-2099), with increases ranging from 28-51% across all studied forests, compared to 10-26% for RCP 2.6 (Table A5). Analysis of long-term trends suggests saturation and subsequent slight decreases in GPP growth rates under RCP 2.6, with this phenomenon being most noticeable in the forests of FI-Hyy. RCP 6.0 consistently shows higher percentage of change in R_eco_ compared to RCP 2.6. By the end of the century (2089-2099), R_eco_ changes from 37-106% for RCP 2.6 and 60-121% for RCP 6.0 across all forests examined (Table A6). Boreal and maritime pine forests experienced higher changes compared to deciduous forests. For example, by 2089-2099, under RCP 6.0, changes reach 142% and 121% at FI-Hyy and FR-Lbr forests, while deciduous forests like European beech in IT-Col and DK-Sor experience lower changes at 67% and 60% respectively.

### 3.3. Changes in NEE dynamics under different climate scenarios

We considered fluctuations in the length of sink/source and source/sink forest stand behaviors throughout the year. This calculation involved determining the total number of days in a year (‘N. days year^—1’^) where a forest stand exhibited either C-sink (NEE < 0) or C-source behavior (NEE > 0). The 10-year average number of days as C-sink and as C-source under the RCP 2.6 and 6.0 for the 5 case studies selected is presented in Figure 5 (panels a). The number of days identified as C-sink in the evergreen forests for the scenario RCP 2.6 (i.e., FR-LBr, CZ-Bk1, and FI-Hyy) starts relatively low in the first decade (1999-2009) but increases significantly over time, showing a consistent upward trend (Figure 5 – 2.a, 3.a and 4.a). At the CZ-Bk1 site, the number of days considered as C-sink starts at 47 N. days year^—1^ in the 1999-2009 decade and increases steadily over the decades, peaking at 199 N. days year^—1^ in 2059-2069 and remains relatively high thereafter. For FI-Hyy forests the number of the days as C-sink starts at 107 N. days year^—1^ in 1999-2009 and rises to 217 days in 2089-2099, with a peak of 225 in 2079-2089. Last, at the forests of FR-LBr, there were no days of C-sink in 1999-2009, but they increased steadily over time, reaching 114 days by 2079-2089. The number of days as C-source varies inversely to the C-sink across decades, demonstrating a general reduction over time for evergreen forests. The number of days functioning as a C-source decreased, from ∼300 in 1999-2009 to ∼200 N. days year^—1^ in 2089-2099 (Table A1), for evergreen forests. Deciduous forests on the contrary almost maintain a relatively stable number of days as C-sink through the century (Figure 5 – 1.a and 5.a). The number of days as C-sink at IT-Col forests ranges from 212 to 224 N. days year^—1^, while at DK-Sor from 214 to 222 N. days year^—1^, with slight fluctuations observed across decades and no clear overall trend. The same trend for the capacity of the stand to act as a C-source, but with a slightly different day in the range from 137 to 159 N. days year^—1^ recorded in the forest of Sorø, and from 140 to 149 N. days year^—1^ for the IT-Col site. The discrepancies between the already described RCP 2.6 and RCP 6.0 scenarios are minimal as they exhibit a similar trend for evergreen and deciduous, respectively, with only a slight change in the number of days, mainly in the last decade, with a magnitude of ∼10 N. days year^—1^.

**Figure 5.**
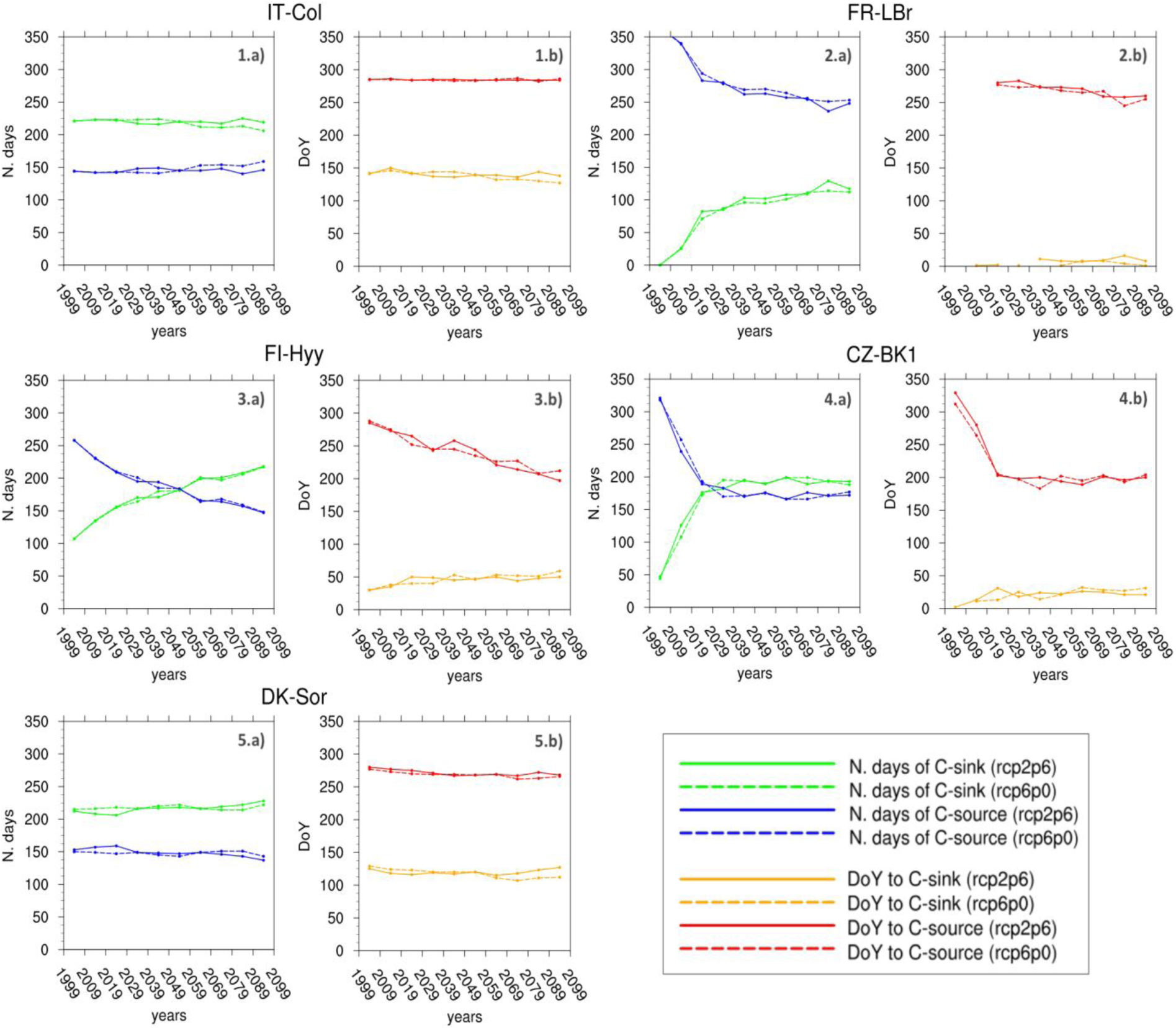
10-year average number of days (N. days year^—1^) as C-sink and as C-source (Figures ‘a’) and DoY (Day of Year) in which C-source switch to C-sink and C-source to C-sink (Figures ‘b’) under the RCP 2.6 and 6.0 for the 5 case studies selected, i.e.: 1) Collelongo - IT-Col, 2) Le Bray - FR-LBr, 3) Hyytiälä - FI-Hyy, 4) Bílý Kříž - CZ-Bk1, and 5) Sorø - DK-Sor). Data missing for some intervals are because of filtering and data removals to avoid pulsing artefacts e.g. ‘Birch effect’ and unstable conditions (see Material and Methods).

To assess the changes in MSC due to climate change scenarios, we focused on the shifting patterns of the DoY over decades, particularly examining when forest stands transitioned from C-sources to C-sink and *vice versa* (see Figure 5, panel b). Regardless of the scenario considered, the European beech forests revealed constant transition periods across decades. In IT-Col, the shift from being a C-source to a C-sink occurred between DoY 136 and 150, with the opposite transition from sink to source around DoY 285. Meanwhile, in DK-Sor, the same transitions happened between DoY 115 and 127, and reversed between DoY 272 and 280 (Figure 5 – 1.b and 5.b). Even for evergreen forests, there are no noticeable differences in the shift corresponding to RCP 2.6 and RCP 6.0 scenarios. At CK-BZ1, the shift to a C-sink occurs relatively early in the year, spanning from DoY 2 to 31 across the decades. At FI-Hyy, the transition timings vary widely, ranging from DoY 30 to 50 over the decades. Similarly, at FR-LBr, the C-sink transitions occur from DoY 1 to 16 across the decades, with some decades exhibiting earlier shifts. On the other hand, transitions to C-source at CZ-BK1 occur from late November to early December (DoY 201-329), displaying a decreasing trend over the years. In contrast, at FI-Hyy, transitions take place from mid-September to late September (DoY 197-285), while at FR-LBr, they occur from late September to early October (DoY 255-285) (Figure 5 – 2.a, 3.a and 4.a).

## 4. Discussion

### 4.1. Model validation

To sum up statistical metrics, the 3D-CMCC-FEM performs best in replicating the mean seasonal patterns of the three fluxes in European beech forests DK-Sor and IT-Col. R_eco_ reaches a satisfactory performance also for the boreal Scots pine forest of FI-Hyy, and the Scots pine forest of CZ-Bk1. The robust predictive ability of 3D-CMCC-FEM in estimating NEE across different timeframes, forest species, and climates, as proved by its alignment with the eddy covariance Fluxnet 2015 Dataset [56], underscores its effectiveness in capturing the complex dynamics of carbon fluxes within forest ecosystems, as documented by previous works (see e.g., [10], [11], [36], [40], [42]). Nevertheless, there are slight inconsistencies, particularly during peak photosynthesis periods (around the 200th day of the year) in both deciduous and evergreen forests, which have been acknowledged in existing literature. Studies indicate that estimates of ecosystem respiration derived from eddy covariance often underestimate actual values for forest ecosystems [72]–[75].

### 4.2. Mean seasonal NEE cycle under climate change scenario

The sensitivity of forest ecosystems to changes in environmental factors such as climate change, seasonal variations, and atmospheric CO_2_ levels has been thoroughly evidenced in the literature. Indeed, the interaction among these variables plays a crucial role in shaping the carbon exchange dynamics [6], [12], [17], [31], [76].

The analysis of the mean NEE seasonal cycle under different climate change scenarios presents several fascinating findings. The first key observation is the consistent reduction of the forest stands capabilities to act as carbon sink from the atmosphere in the coming years, across all study sites, and climate change scenarios. Despite the overall lack of understanding regarding how forest ecosystems will respond to climate change, several new studies concur regarding the diminishing forests carbon uptake capabilities, thus confirming our statement [26], [31], [77]– [79].

In broader terms, climate change affects forest carbon balance by influencing key processes, which can respond differently due to their sensitivity to various environmental drivers [80]. Indeed, the reduction in NEE, within the forests observed in this study, has a disparity in intensity of climate impacts across these scenarios: RCP 2.6 exhibits less intense NEE reduction compared to RCP 6.0 scenarios. The warming accelerates both autotrophic and heterotrophic respiration meaning that increased temperature may lead forests to an increase of R_eco_ [27], [58], [81], [82]. On the other hand, the increasing atmospheric CO_2_ concentration intensifies the GPP through the ‘carbon fertilization effect’ (i.e., reported to be the cause of 44% GPP increase since the 2000s) [18], [19]. Under the RCP 2.6 scenario, R_eco_ exhibits a steady linear increase until the century’s end, while GPP follows a bell-shaped curve, reaching saturation around the mid-century. Consequently, as GPP saturates, its compensatory capacity reduces, while R_eco_ continues to rise due to further temperature increases in the latter half of the century resulting in a further decline in NEE. Conversely, for the RCP 6.0 scenario, GPP saturation occurs only towards the end of the century, and while one might anticipate a reversal in NEE trends considering this factor, it does not materialize. This is because under the RCP 6.0 scenario, a higher temperature increase is predicted compared to RCP 2.6 (i.e., [58]), leading to a more pronounced rise in ecosystem respiration relative to photosynthesis. As a consequence, there’s a greater decrease in the stand forest’s carbon sink capacity. The result refers to all forests studied in this work, but in the forest of FI-Hyy the phenomena look more pronounced (Figures A2 and A3 in Appendix). The fate of ecosystem carbon flux depends, not only on atmospheric and climate conditions but also on the type of forests analyzed. The reduction of C-sink capabilities is particularly notable in evergreen forests, which exhibit a higher decrease in NEE compared to evergreen sites. The boreal Scots pine forest of FI-Hyy stands out with the most significant reduction in NEE, indicating a heightened vulnerability to climate change effects. This finding is supported by the study of Hadden and Grelle [26] who found, that over a 17-year period, the forest ecosystem in a boreal forest stand in northern Sweden transitioned from being a carbon sink to a carbon source. This could mean that the past efforts to validate the neutrality hypothesis [83] with climate change impact showing limitations, and we need new research directions and new perspectives to better capture changes in the carbon fluxes between the ecosystem and atmosphere [84]. Indeed, a long-lasting debate around the C-neutrality of old-growth forests (and some of the sites become old-growth at the end of simulations) raised concerns and increasing debates about the sink capacity at forest ageing as Odum’s theory would assume. As found in [7] (but see [85]) we also found that even a >200-year-old stand (as IT-Col in 2099) still has sink capacity. The annual NEE decrease (and much less the changes in the MSC) as shown by the model results is certainly a function not only of climate but also of the inherent effects related to e.g., biomass, both live and dead, accumulation (which led to increases in respiratory costs) and changes in e.g., the forest structure (which led to decreases in the carbon assimilation) as stands age. Nevertheless, such effects on annual NEE are expected to be greatly exacerbated by climate change.

### 4.3. Changes in NEE dynamics under different climate scenarios

Climate change impacts plant phenology by altering the start and end dates of the growing season, which influences when photosynthesis can begin and consequently affects C-fluxes [20], [29], [30], [32], [86].

A primary finding is that, regardless of the scenario analyzed, the number of days identified as C-sink in evergreen forests increases significantly over time, indicating a consistent upward trend. Similarly, the number of days classified as C-source decreases over the decades, showing a general reduction. The second finding is that for evergreen forests, the DoY to C-sink tends to increase (indicating a forward shift in the year when the system becomes a sink), and the DoY to C-source decreases (indicating a backward shift in the year when the system becomes a source), aligning with the overall trend of fewer C-sink days and more C-source days over time. In contrast, deciduous forests maintain a relatively stable number of C-sink (and C-source) days throughout the century, reflecting a steady DoY when the system becomes a C-sink (or C-source), despite an anticipated beginning of the growing season but compensated by higher respiration rates. Indeed, for the deciduous, the 3D-CMCC-FEM simulates the bud-brake through a thermic sum function and leaf and fine root development (and the relative growth respiration peak) in a well-defined and short time during spring [36]. Conversely, for the evergreens leaf and fine root growth development is spread on all over the spring. At the same time, leaf fall in the deciduous starts under certain hours of solar radiation, and thus, this is not under the control of climate, while in the evergreen it happens all over the year, and under the control of climate, balanced by incoming photosynthates for new leaves and fine roots. Ultimately, deciduous spring C-sink capacity is counterbalanced by high C-emissions because of mainly growth respiration in spring. Such behavior is different for evergreens, which length their C-sink capacity during spring. However, the lengthening of the growing season does not automatically mirror an increase in the net sink capacity because R_eco_ shows to increase much more than GPP. It is generally acknowledged that the changing temperature response of respiration transforms forests from C-sinks to C-sources [26], [76], [87]–[89], while the stability of the carbon sink/source dynamics over the decades for deciduous ecosystems is a relatively recent finding. The DoY for deciduous forests remains unchanged because the earlier start of the growing season, triggered by rising temperatures, is balanced by an earlier increase in respiration. This compensates for the earlier rise in GPP at the level of NEE. Overall, the lack of change in the number of C-sink (and C-source) days across decades and the reduction of the NEE suggest that, over the long term, deciduous forests are more efficient in using photosynthates compared to evergreen forests [90], [91].

### 4.4. Limitations

The modeling framework presented has certain limitations that must be acknowledged. First, we deliberately decide to not simulate neither the effects of anthropogenic disturbances, e.g., forest management, not the ones from natural disturbances caused by climate change, such as windstorms, forest fires, and insect outbreaks to concentrate on the effects of climate change alone and avoiding these potentially confounding effects Climate extreme events are presumed to be incorporated in the climate scenarios used to drive the model and then, somewhat, already accounted for. Moreover, indirect changes due to climate change in key factors like nitrogen deposition, phosphorus, or ozone—which could potentially amplify or mitigate our findings— were not evaluated. Nonetheless, some research (e.g., [92]) indicates that this issue might not be universally relevant. These studies highlight the strong response of various tree species to CO_2_ fertilization across different levels of nutrient availability. Lastly, the potential for species migration to and from the study areas was not considered. However, such dynamics might require longer timescales than those covered in this study and it is unlikely (although still possible in theory) that species composition may completely change throughout the simulations.

## 5. Conclusion

The MSC metric is an interesting and descriptive metric that associates phenology and carbon partitioning allocation within forest stands. Climate change impacts both the phenology by changing the date for the beginning and the end of the growing season, and the ecosystem carbon allocation.

We applied the process-based forest model 3D-CMCC-FEM to evaluate the potential modifications in the ability of different forest stands to absorb or emit carbon throughout the season and across the years up to 2100. Before that, we validate the model under current climate conditions and find a robust predictive ability of 3D-CMCC-FEM in estimating NEE, GPP and R_eco_ across different timeframes, forest species, and climates.

The analysis of the mean NEE seasonal cycle under different climate change scenarios presents the consistent reduction of the forest stands capabilities to act as carbon sink in the coming years, across all study sites, and climate change scenarios. The reduction in NEE ability has different intensities of climate impacts across these scenarios. The RCP 2.6 scenario demonstrates a less pronounced reduction NEE compared to the RCP 6.0 scenario. This disparity primarily stems from variations in key variables, such as the differing rates of temperature increase between the two scenarios, as well as the CO_2_ fertilization effect while in all sites age effects depend on the age at the beginning of simulations. The reduction of C-sink capabilities is mainly notable in evergreen forests, which exhibit a higher decrease in NEE compared to deciduous forest sites. Finally, we found that the number of days as C-sink in evergreen forests increases over the years, indicating a consistent upward trend. Oppositely, the number of days of C-source decreases over the decades, showing a general reduction. This statement aligns with the forward shift of DoY to C-sink, and the backward shift of DoY to C-source. In contrast, deciduous forests maintain a relatively stable number of C-sink (and C-source) days throughout the century, reflecting a fixed DoY when the system becomes a C-sink (or C-source). The DoY for deciduous forests remains constant as the earlier onset of the growing season, driven by warming temperatures, is offset by an earlier uptick in respiration. Decades pass with little change in the number of days as C-sink (and C-source), alongside a decrease in NEE. This indicates that deciduous forests, over the long haul, demonstrate greater efficiency in utilizing photosynthates when compared to evergreen forests.

## Author Contributions

M. Morichetti: Data curation, Formal analysis, Investigation, Writing - original draft, Writing - review & editing; E. Vangi: Data curation, Writing – original draft, Writing - review & editing; A. Collalti: Conceptualization, Formal analysis, Investigation, Writing - original draft, Writing - review & editing

## Data Availability Statement

The 3D-CMCC-FEM model code is publicly available and can be found on the GitHub plat-form at: https://github.com/Forest-Modelling-Lab/3D-CMCC-FEM. The 3D-CMCC-FEM output data used in this work can be downloaded at: https://zenodo.org/records/11124413. Correspondence and requests for additional materials should be addressed to the corresponding author.

## Acknowledgements

We are thankful to D. Dalmonech for supporting data preparation and analysis. M.M. and A.C. acknowledge funding by the project OptForEU Horizon Europe research and innovation programme under grant agreement No. 101060554. A.C. also acknowledges the project funded under the National Recovery and Resilience Plan (NRRP), Mission 4 Component 2 Investment 1.4 - Call for tender No. 3138 of 16 December 2021, rectified by Decree n.3175 of 18 December 2021 of Italian Ministry of University and Research funded by the European Union – NextGenerationEU under award Number: Project code CN_00000033, Concession Decree No. 1034 of 17 June 2022 adopted by the Italian Ministry of University and Research, CUP B83C22002930006, Project title “National Biodiversity Future Centre - NBFC”. A.C. also acknowledge funding from the MIUR Project (PRIN 2020) - Research Projects of National Relevance funded by the Italian Ministry of University and Research entitled: “Unraveling interactions between WATER and carbon cycles during drought and their impact on water resources and forest and grassland ecosySTEMs in the Mediterranean climate” (WATERSTEM, project num­ber: 20202WF53Z), and “WAFER” at CNR (Consiglio Nazionale delle Ricerche). A.C. and E.V. also acknowledge funding from the MIUR Project (PRIN 2020) - Research Projects of National Relevance funded by the Italian Ministry of University and Research entitled: “Multi-scale observations to predict Forest response to pollution and climate change” (MULTIFOR, project number: 2020E52THS). We also thank the ISIMIP project (https://www.is imip.org/) and the COST Action FP1304 PROFOUND (Towards Robust Projections of European Forests under Climate Change), supported by COST (European Cooperation in Science and Technology) for providing us the climate historical and future scenarios and site data used in this work. This work used eddy covariance data acquired and shared by the FLUXNET community, including these networks: AmeriFlux, AfriFlux, AsiaFlux, CarboAfrica, CarboEurope-IP, CarboItaly, CarboMont, ChinaFlux, Fluxnet-Canada, GreenGrass, ICOS, KoFlux, LBA, NECC, OzFlux-TERN, TCOS-Siberia, and USCCC. The ERA-Interim reanalysis data are pro-vided by ECMWF and processed by LSCE. The FLUXNET eddy covari-ance data processing and harmonization was carried out by the European Fluxes Database Cluster, AmeriFlux Management Project, and Fluxdata Project of FLUXNET, with the support of CDIAC and ICOS Ecosystem Thematic Center, and the OzFlux, ChinaFlux, and AsiaFlux offices. We acknowledge the World Climate Research Programme’s Working Group on Coupled Modelling, which is responsible for CMIP, and we thank the respective climate modelling groups for producing and making available their model output. The U.S. Department of Energy’s Program for Climate Model Diagnosis and Intercomparison at Lawrence Livermore National Laboratory provides coordinating support for CMIP and led the development of software infrastructure in partnership with the Global Organization for Earth System Science Portals.

## Appendix

**Table A1.**
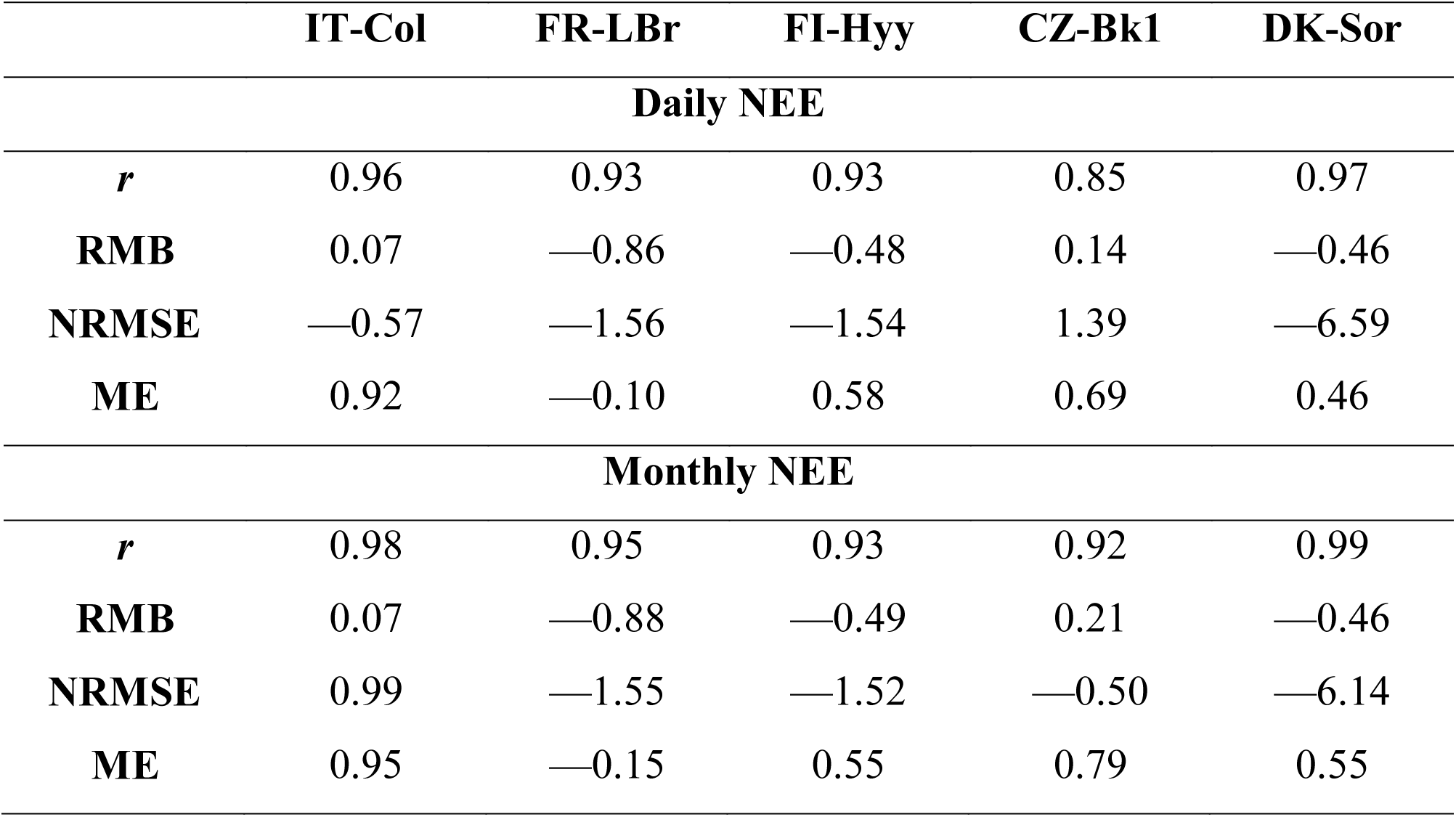
Summary of the statistics between simulated and measured NEE from the Fluxnet2015 Dataset (Pastorello et al., 2020), calculated on the 5 cases studies selected (i.e., Collelongo - IT-Col, Le Bray - FR-LBr, Hyytiälä - FI-Hyy, Bílý Křìž - CZ-Bk1, and Sorø - DK-Sor). The table shows the daily, monthly and yearly values for Person’s Coefficient (r - dimensionless), Relative Mean Bias (RMB - %), Normalized Root Mean Square Error (NRMSE - dimensionless), Modeling Efficiency (ME - %).

**Table A2.**
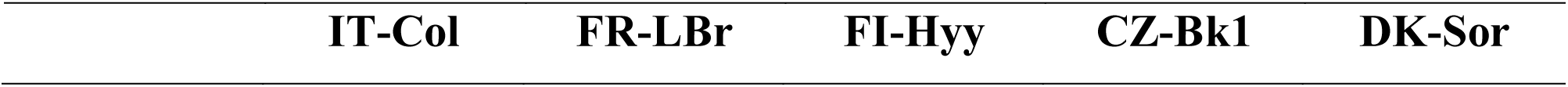
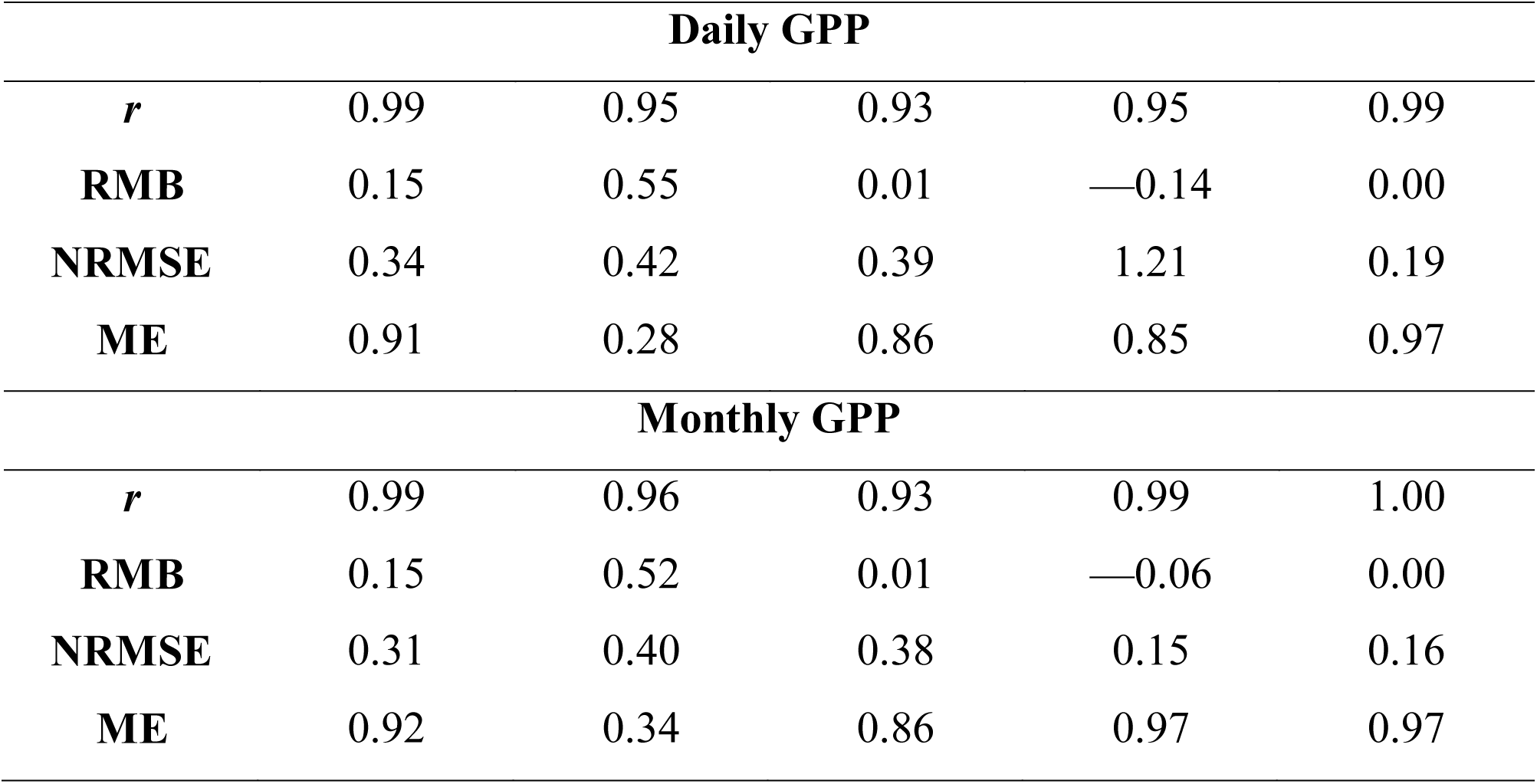
Summary of the statistics between simulated and measured GPP from the Fluxnet2015 Dataset (Pastorello et al., 2020), calculated on the 5 cases studies selected (i.e., Collelongo - IT-Col, Le Bray - FR-Lbr, Hyytiälä - FI-Hyy, Bílý Křìž - CZ-Bk1, and Sorø - DK-Sor). The table shows the daily, monthly and yearly values for Person’s Coefficient (r - dimensionless), Relative Mean Bias (RMB - %), Normalized Root Mean Square Error (NRMSE - dimensionless), Modeling Efficiency (ME - %).

**Table A3.**
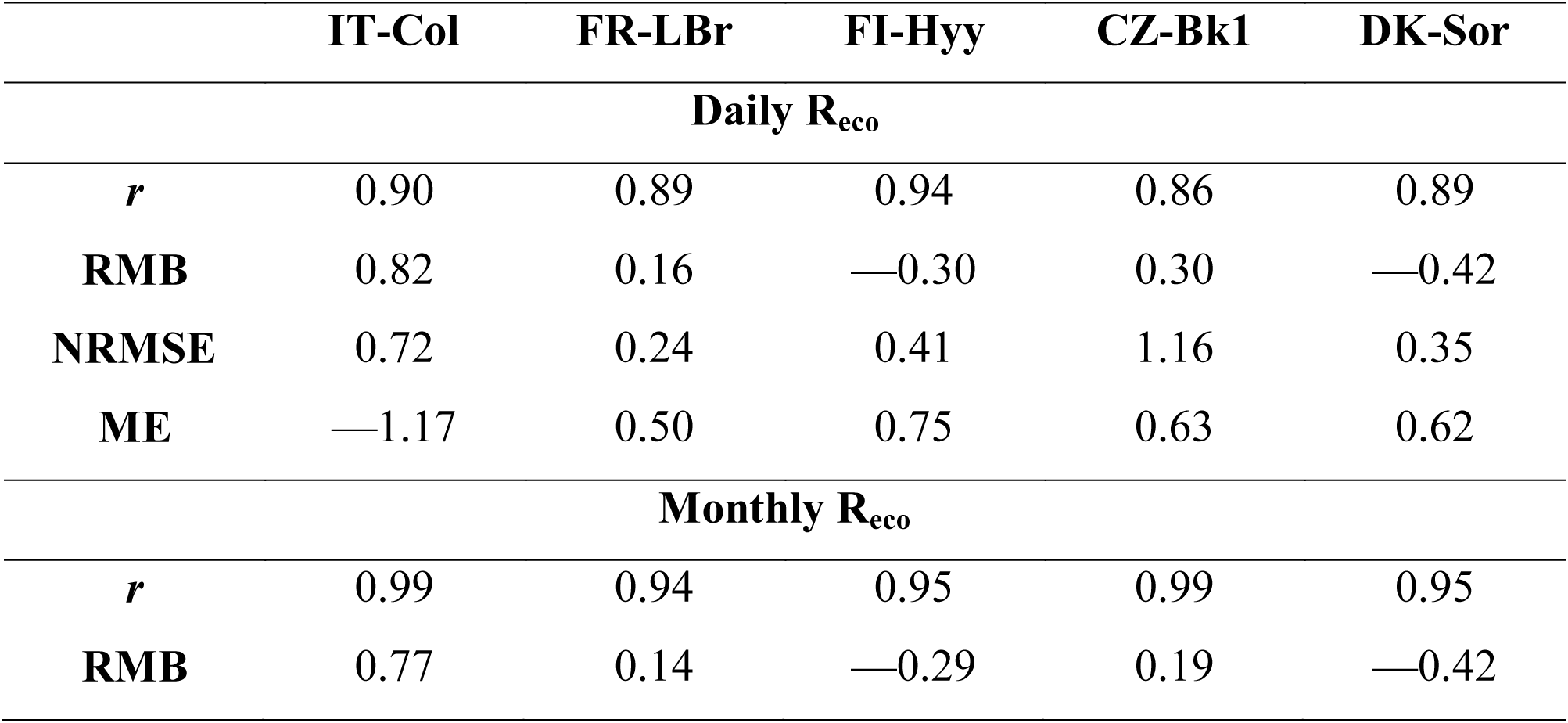
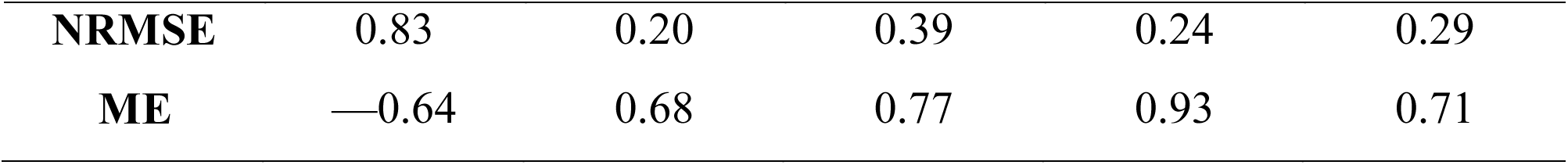
Summary of the statistics between simulated and measured R_eco_ from the Fluxnet2015 Dataset (Pastorello et al., 2020), calculated on the 5 cases studies selected (i.e., Collelongo - IT- Col, Le Bray - FR-LBr, Hyytiälä - FI-Hyy, Bílý Křìž - CZ-Bk1, and Sorø - DK-Sor). The table shows the daily, monthly and yearly values for Person’s Coefficient (r - dimensionless), Relative Mean Bias (RMB - %), Normalized Root Mean Square Error (NRMSE - dimensionless), Modeling Efficiency (ME - %).

**Table A4.**
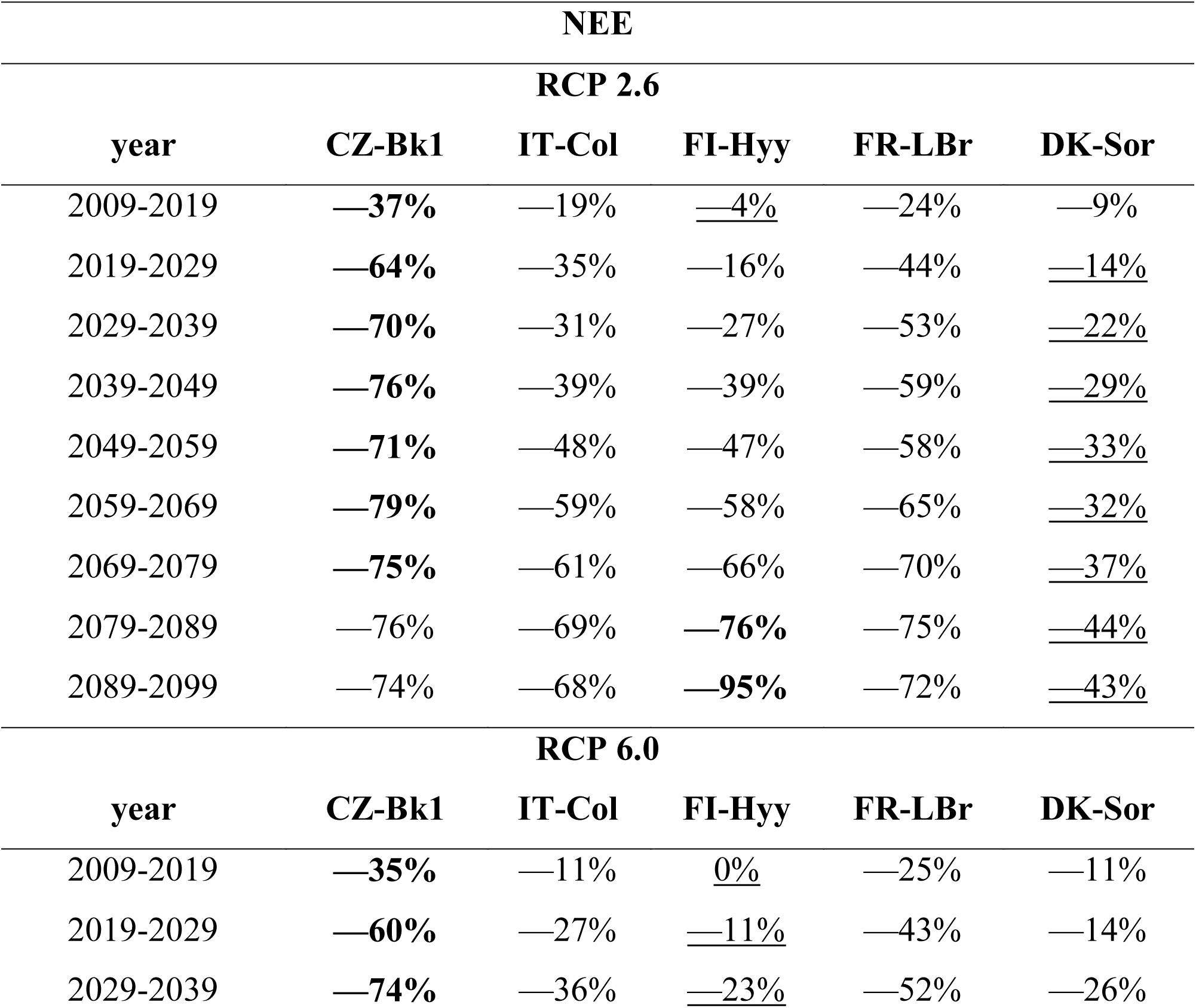
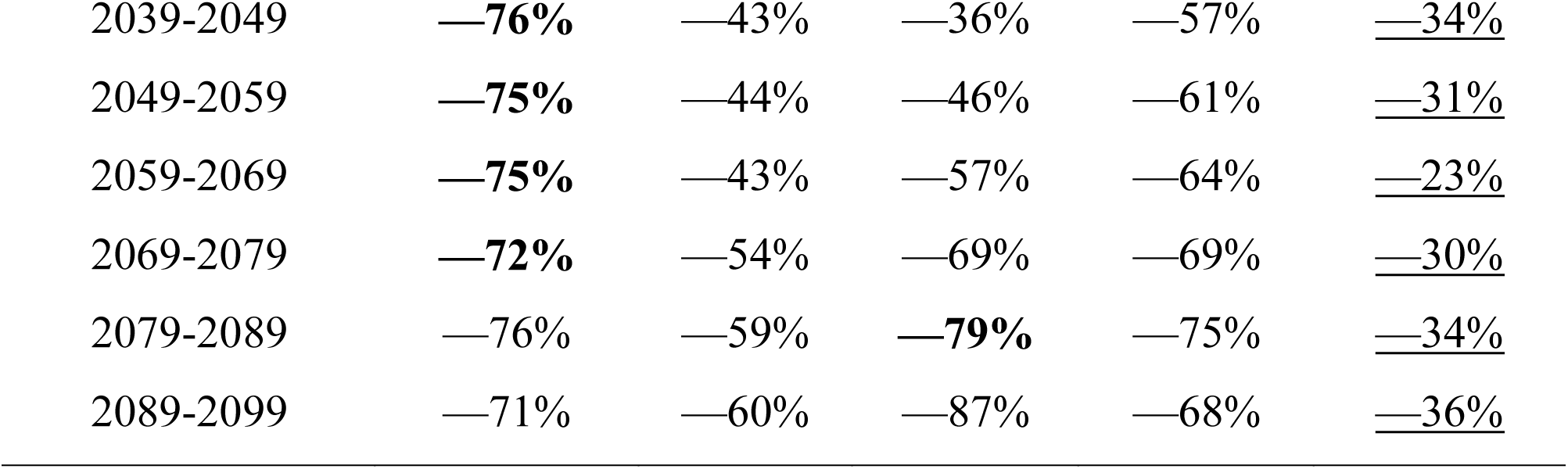
Summary of the NEE variation (%) from the first decade of simulation (1999-2009), considering both climate change scenarios (RCP 2.6 and 6.0) for 5 case studies selected (i.e., Collelongo - IT-Col, Le Bray - FR-LBr, Hyytiälä - FI-Hyy, Bílý Kříž - CZ-Bk1, and Sorø - DK-Sor). In bold values where changes were the highest between the decades while underlined the lowest ones. Note that negative values indicate that NEE becomes less negative, e.g. —100% indicates a reduction in the negative values of NEE.

**Table A5.**
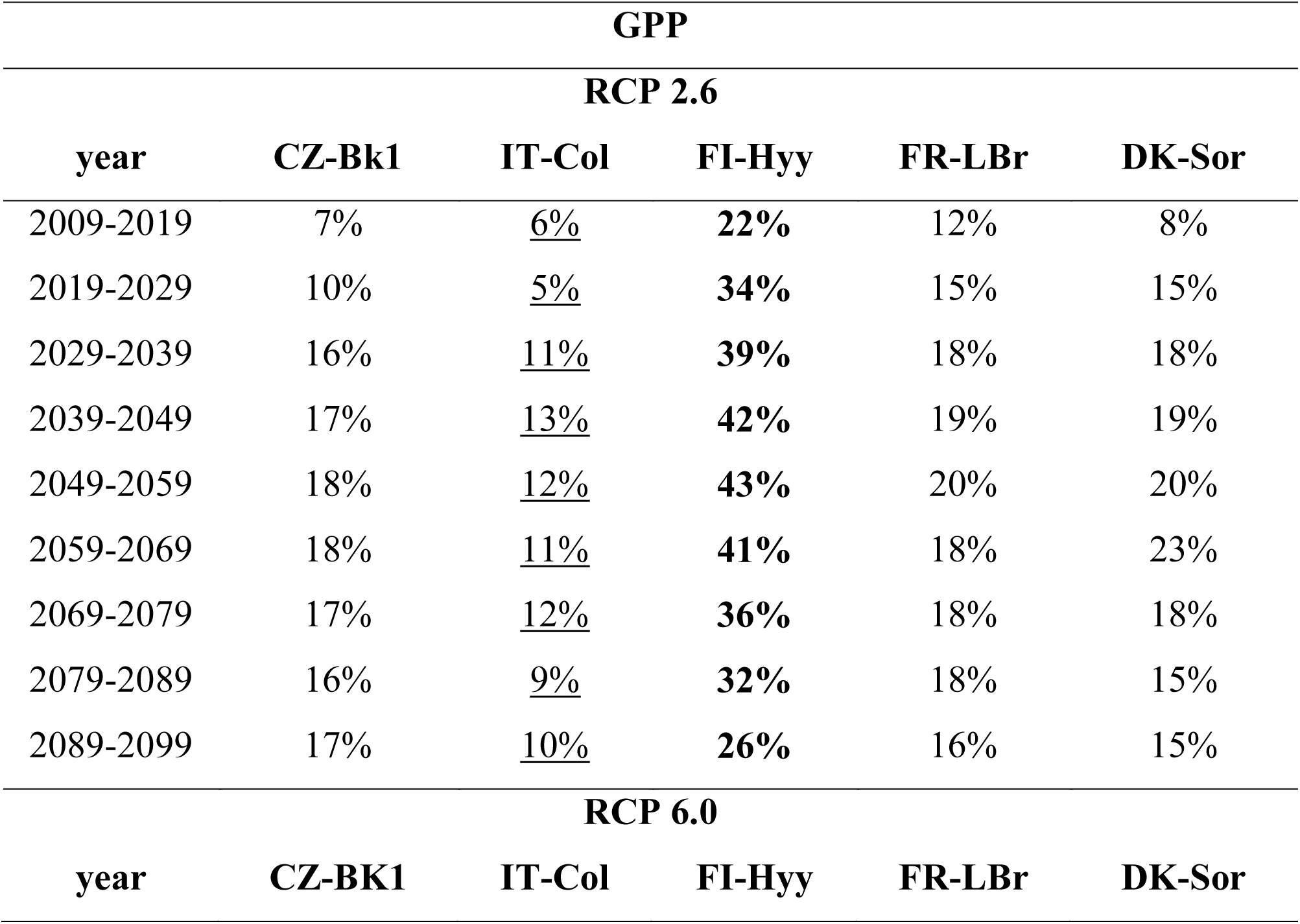
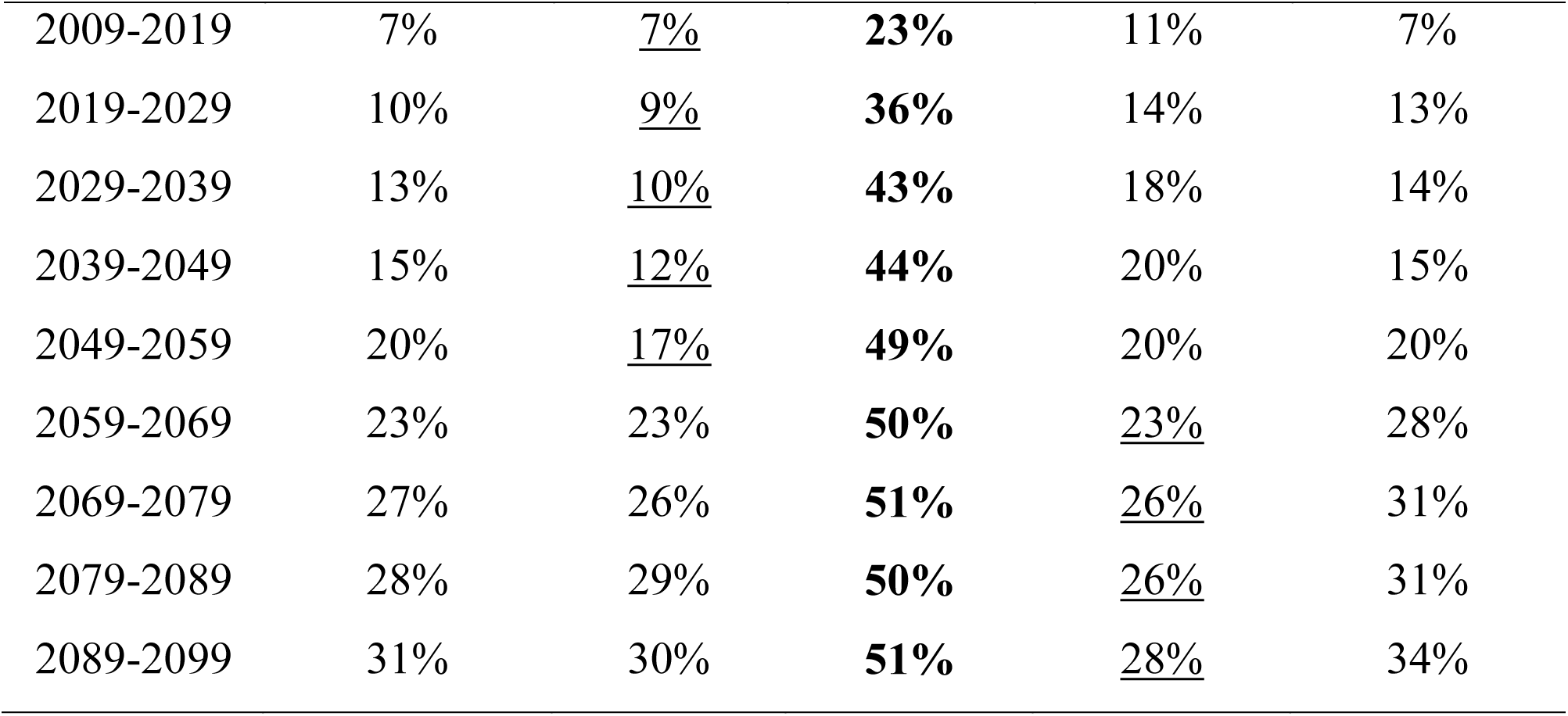
Summary of the GPP variation (%) from the first decade of simulation (1999-2009), considering both climate change scenarios (RCP 2.6 and 6.0) for 5 case studies selected (i.e., Collelongo - IT-Col, Le Bray - FR-LBr, Hyytiälä - FI-Hyy, Bílý Kříž - CZ-Bk1, and Sorø - DK-Sor). In bold values where changes were the highest between the decades while underlined the lowest ones.

**Table A5.**
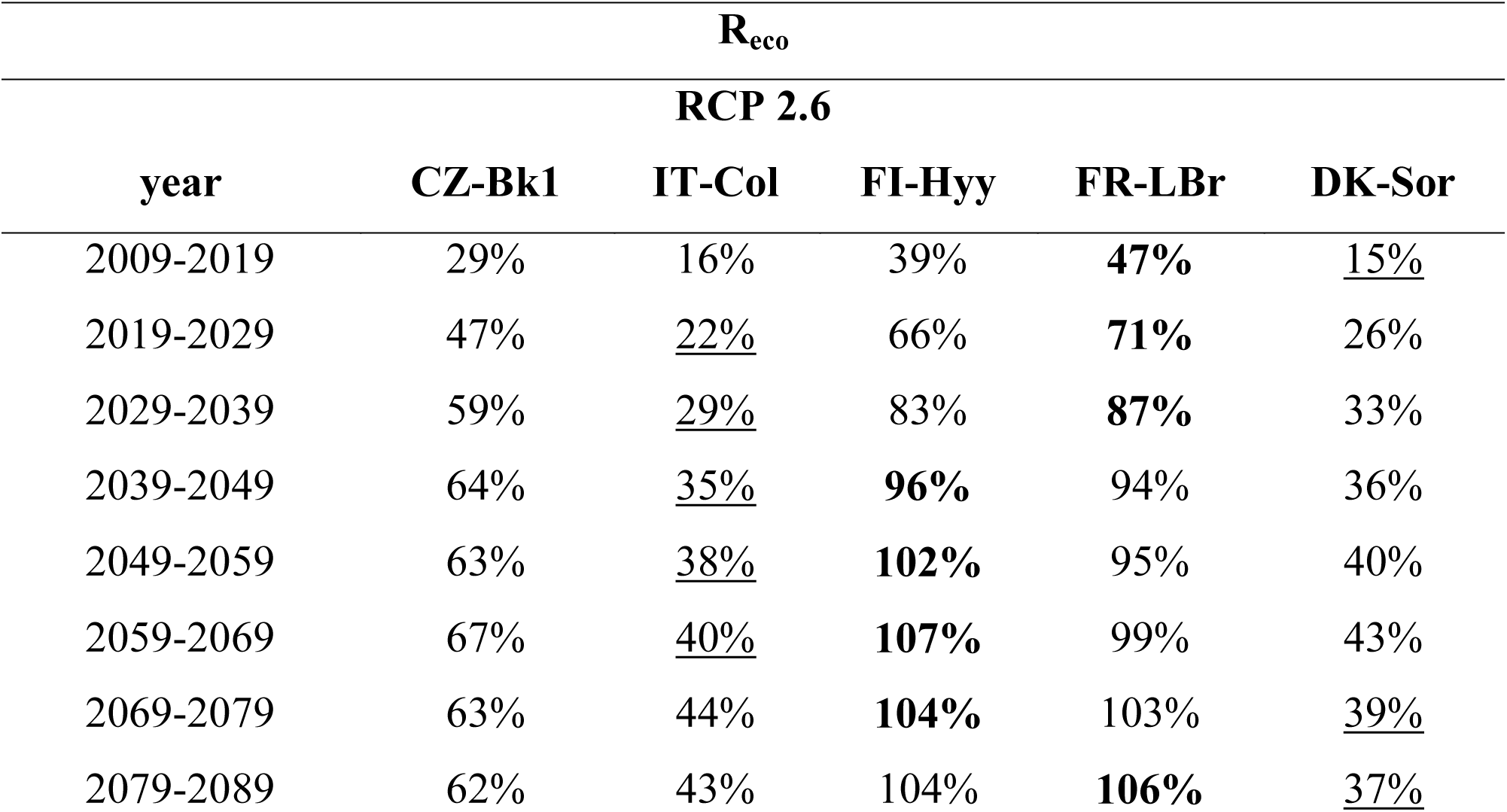
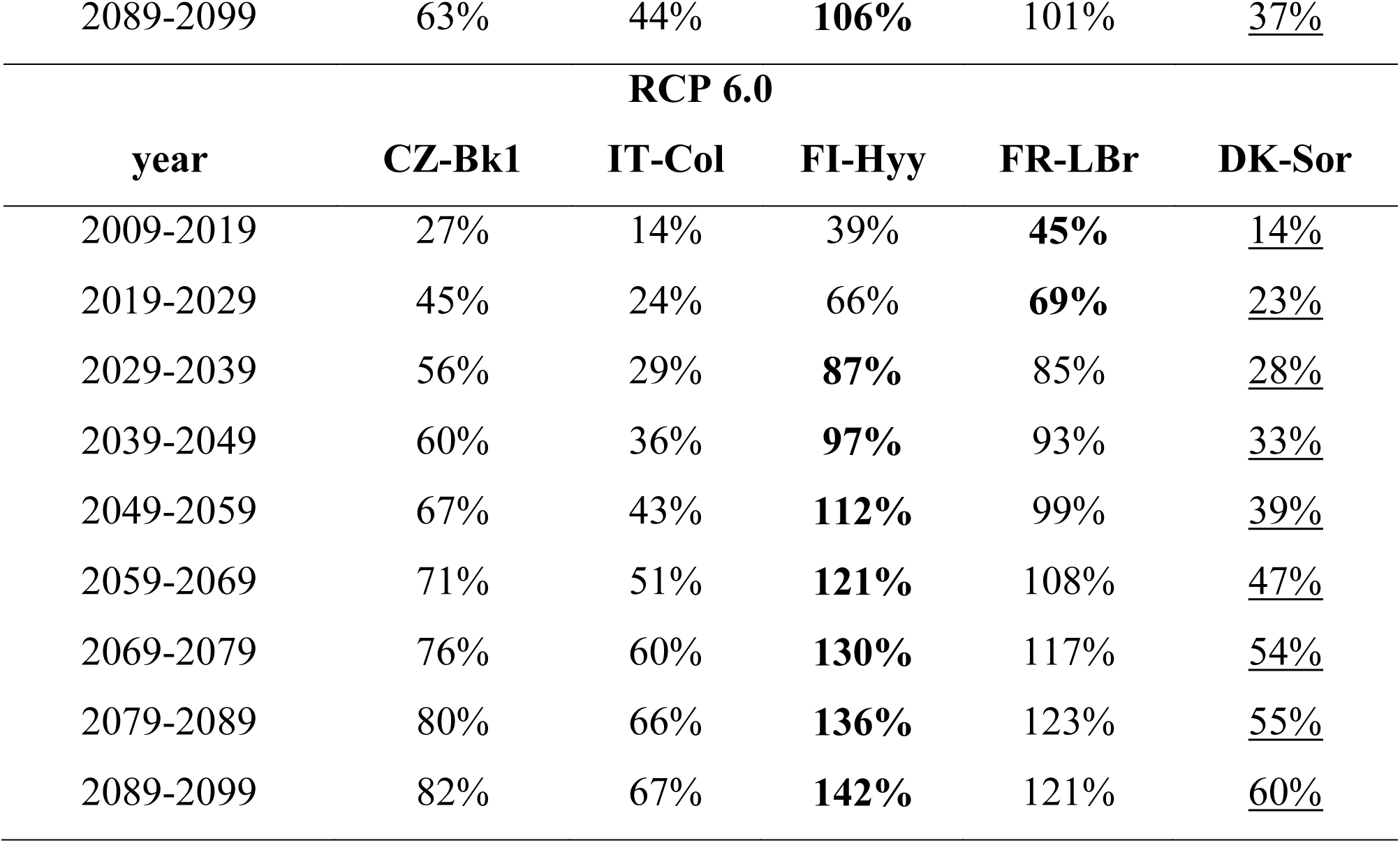
Summary of the R_eco_ variation (%) from the first decade of simulation (1999-2009), considering both climate change scenarios (RCP 2.6 and 6.0) for 5 case studies selected (i.e., Collelongo - IT-Col, Le Bray - FR-LBr, Hyytiälä - FI-Hyy, Bílý Kříž - CZ-Bk1, and Sorø - DK-Sor). In bold values where changes were the highest between the decades while underlined the lowest ones.

**Table A6.**
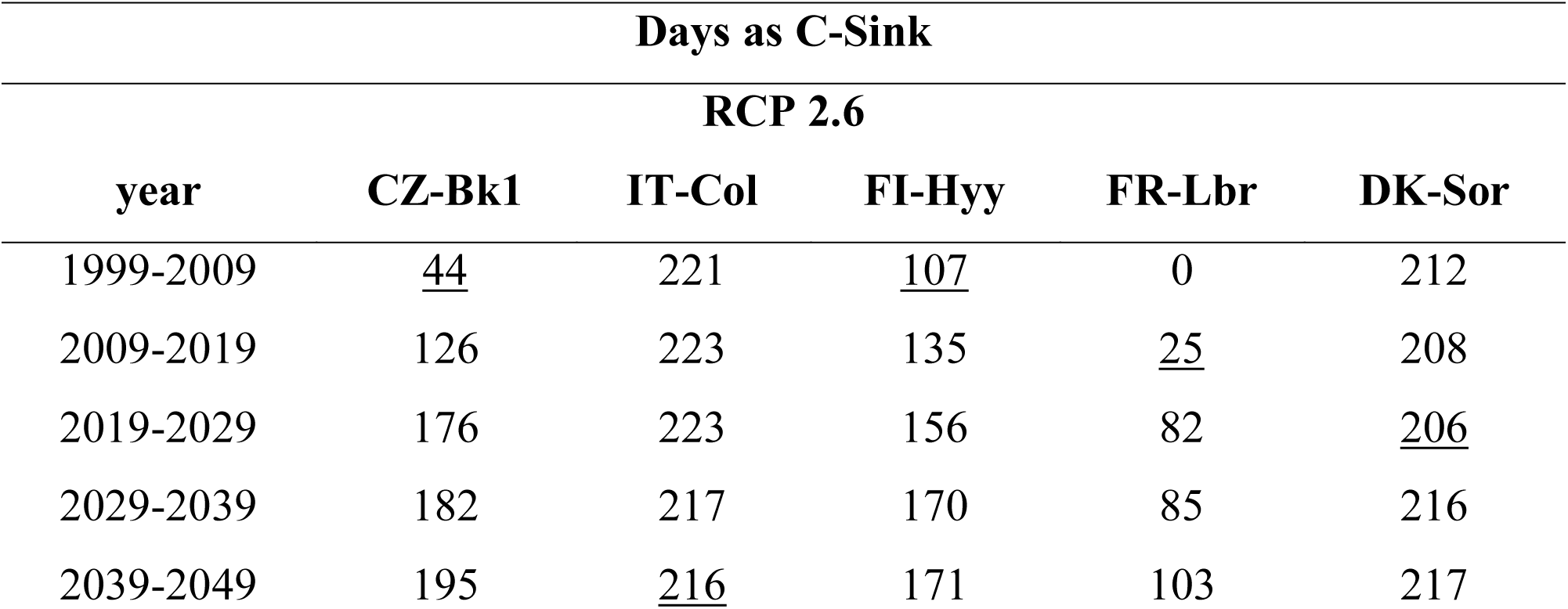
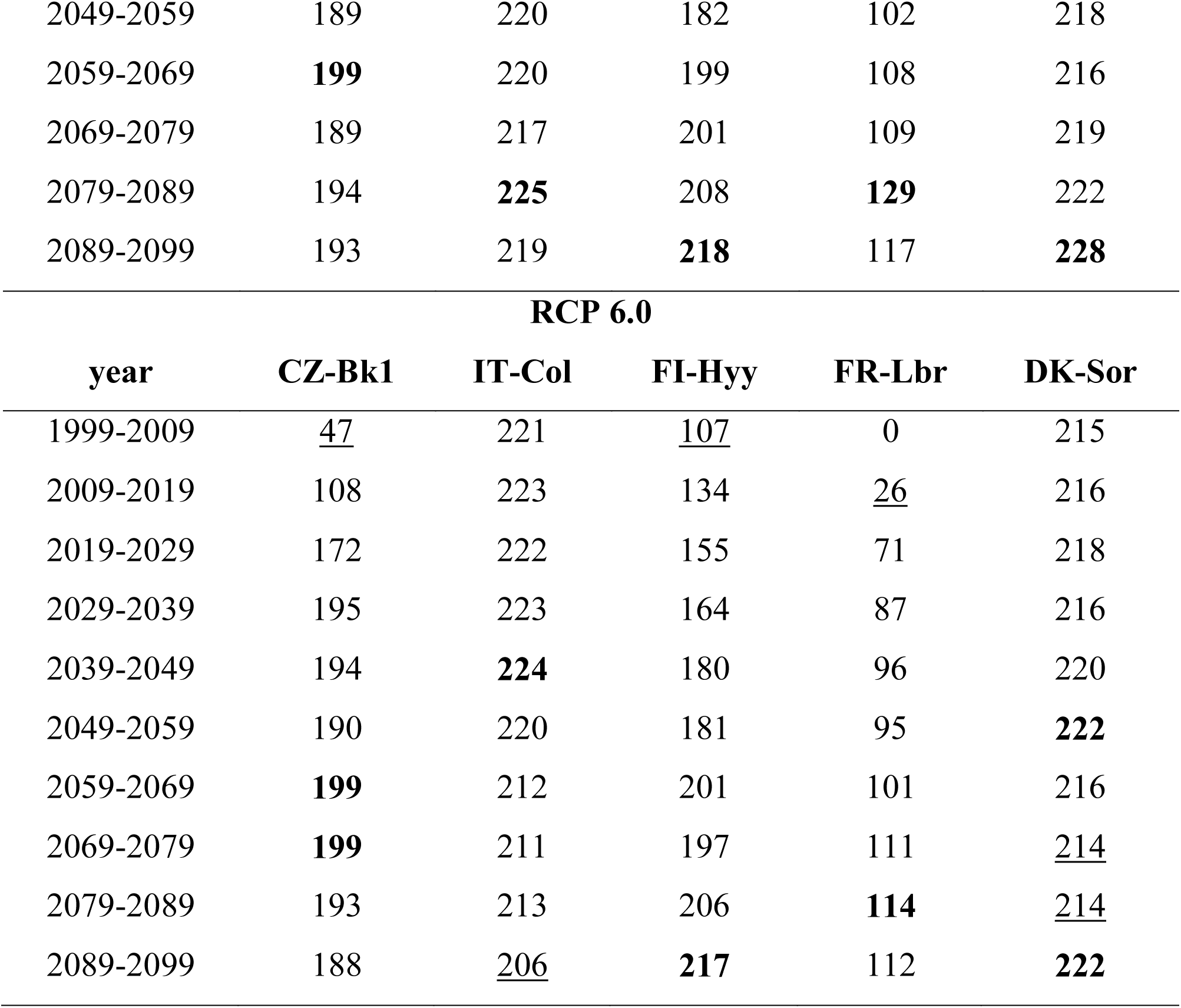
Summary of the number of the days in a year (N. days year^−1^) as C-sink, considering both climate change scenarios (RCP 2.6 and 6.0) for 5 case studies selected (i.e., Collelongo - IT-Col, Le Bray - FR-LBr, Hyytiälä - FI-Hyy, Bílý Kříž - CZ-Bk1, and Sorø - DK-Sor). In bold values where changes were the highest between the decades while underlined the lowest ones within each forest stand.

**Table A7.**
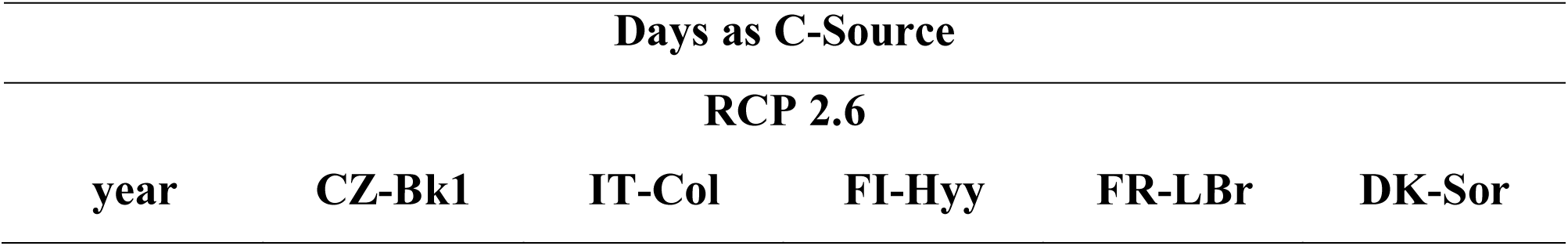
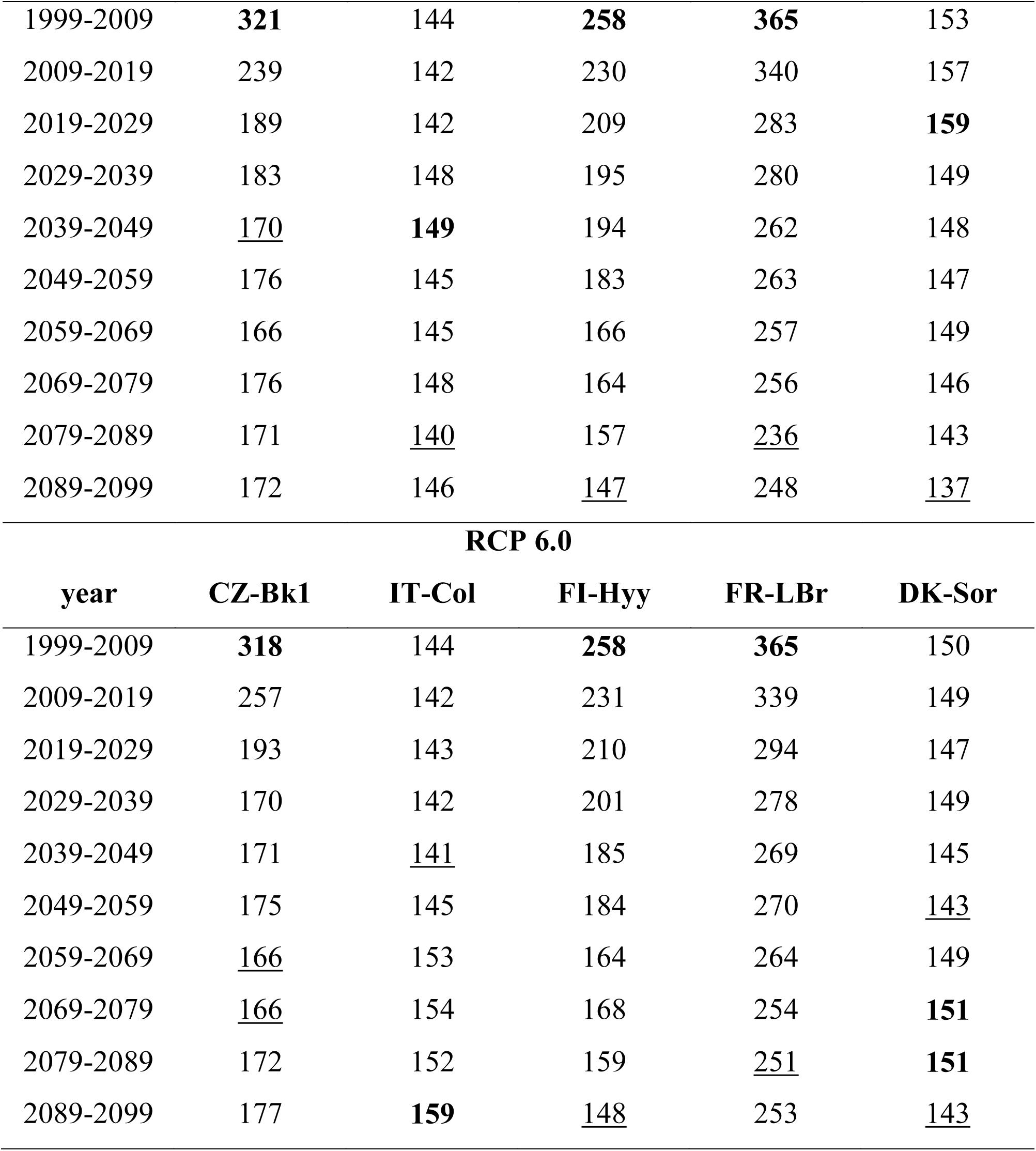
Summary of number of the days in a year (n. days year^−1^) source, considering both climate change scenarios (RCP 2.6 and 6.0) for 5 case studies selected (i.e., Collelongo - IT-Col, Le Bray - FR-LBr, Hyytiälä - FI-Hyy, Bílý Kříž - CZ-Bk1, and Sorø - DK-Sor). In bold values where changes were the highest between the decades while underlined the lowest ones within each forest stand.

**Table A8.**
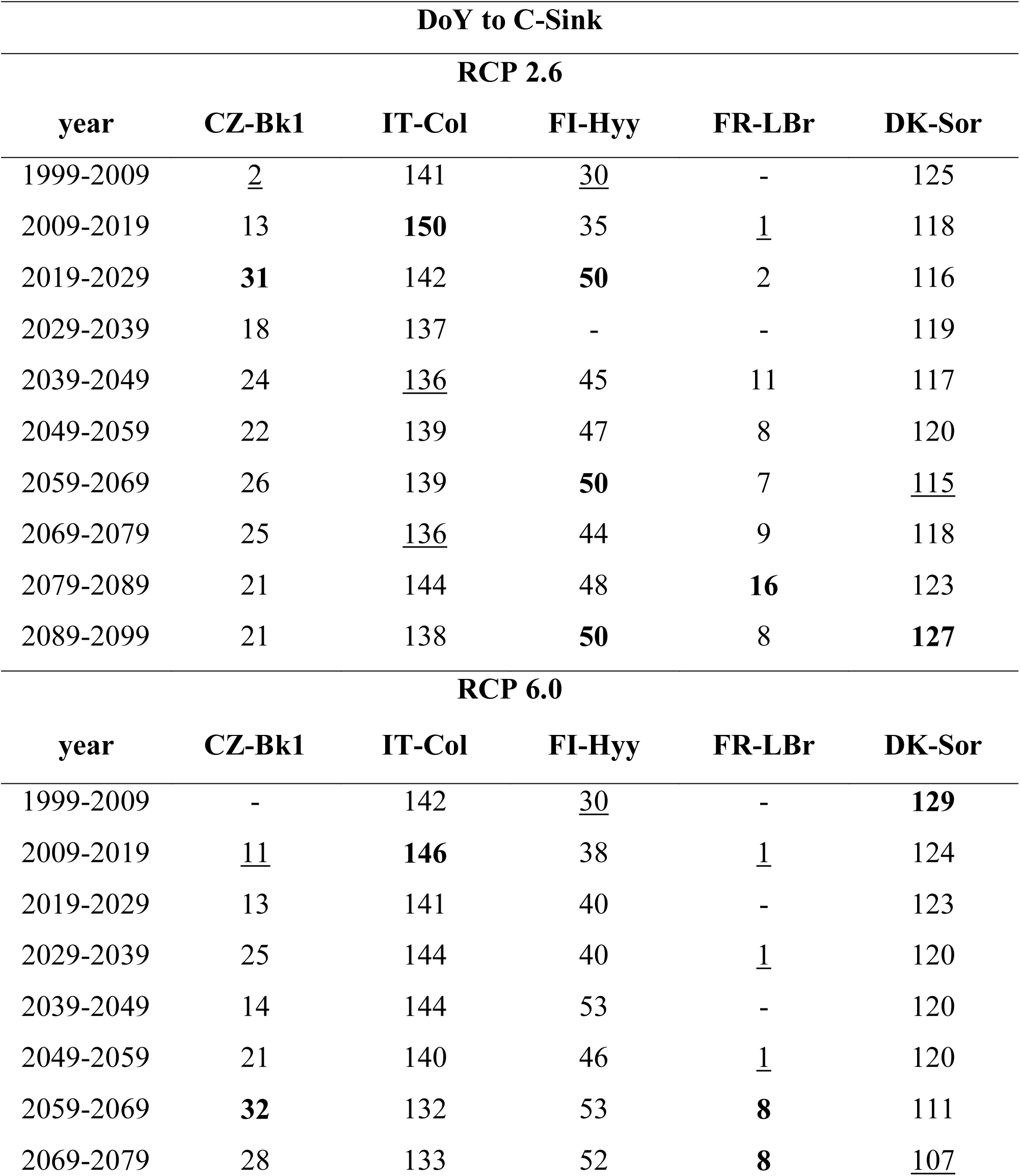
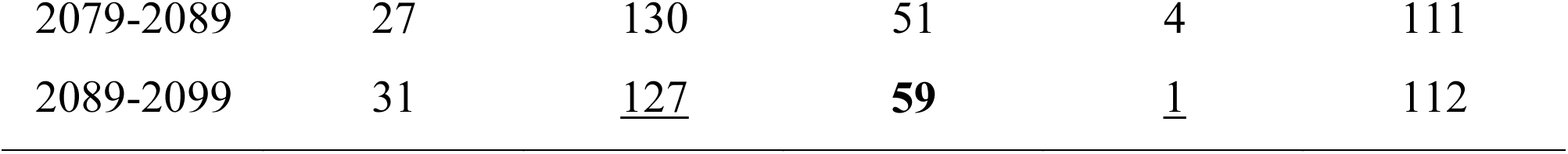
Summary of the changes in the source/sink DoY (Day of Year), considering both climate change scenarios (RCP 2.6 and 6.0) for 5 case studies selected (i.e., Collelongo - IT-Col, Le Bray - FR-LBr, Hyytiälä - FI-Hyy, Bílý Kříž - CZ-Bk1, and Sorø - DK-Sor). In bold values where changes were the highest between the decades while underlined the lowest ones within each forest stand. Data missing for some intervals are because of filtering and data removals to avoid pulsing artefacts e.g. ‘Birch effect’ (see Material and Methods).

**Table A9.**
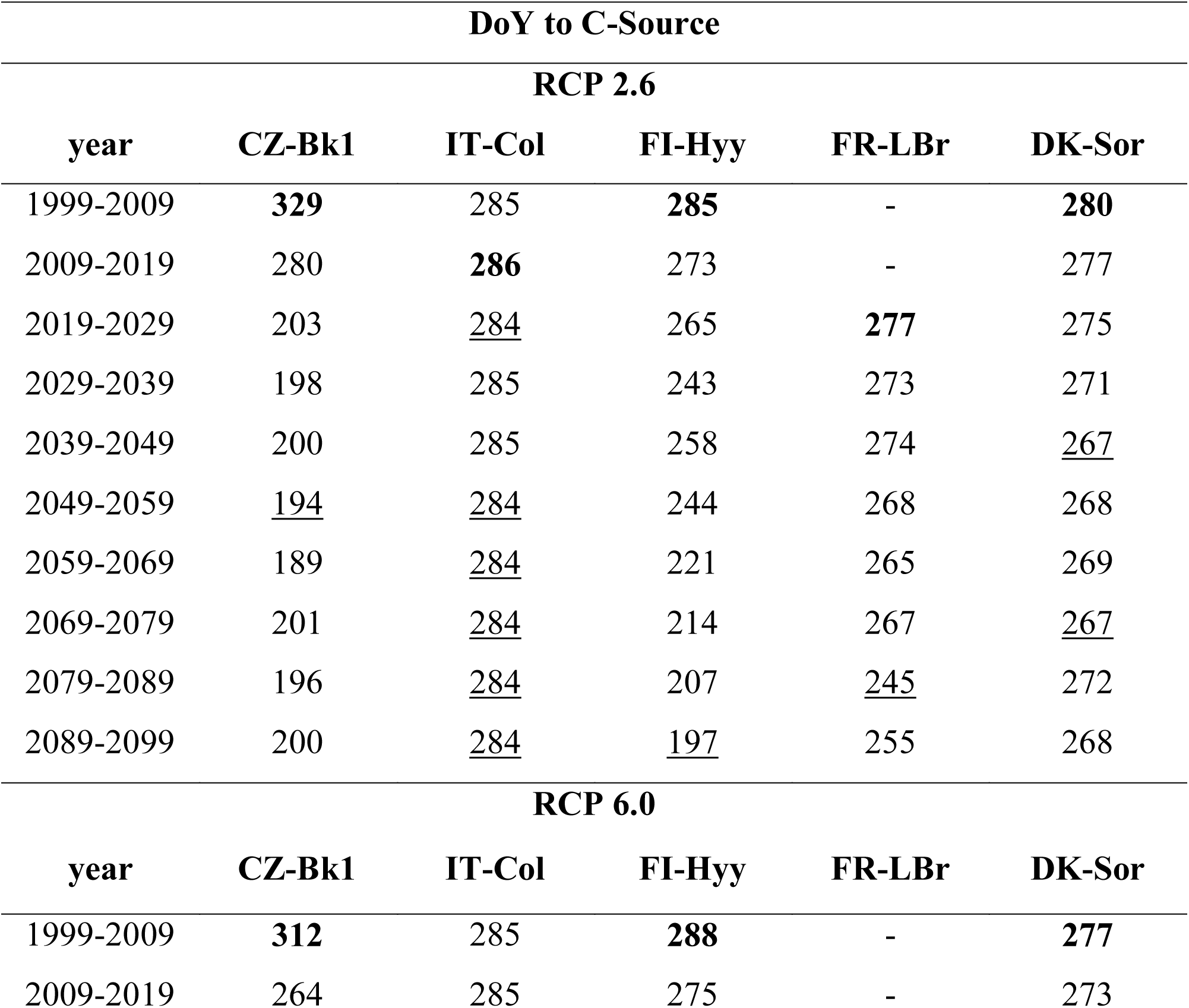
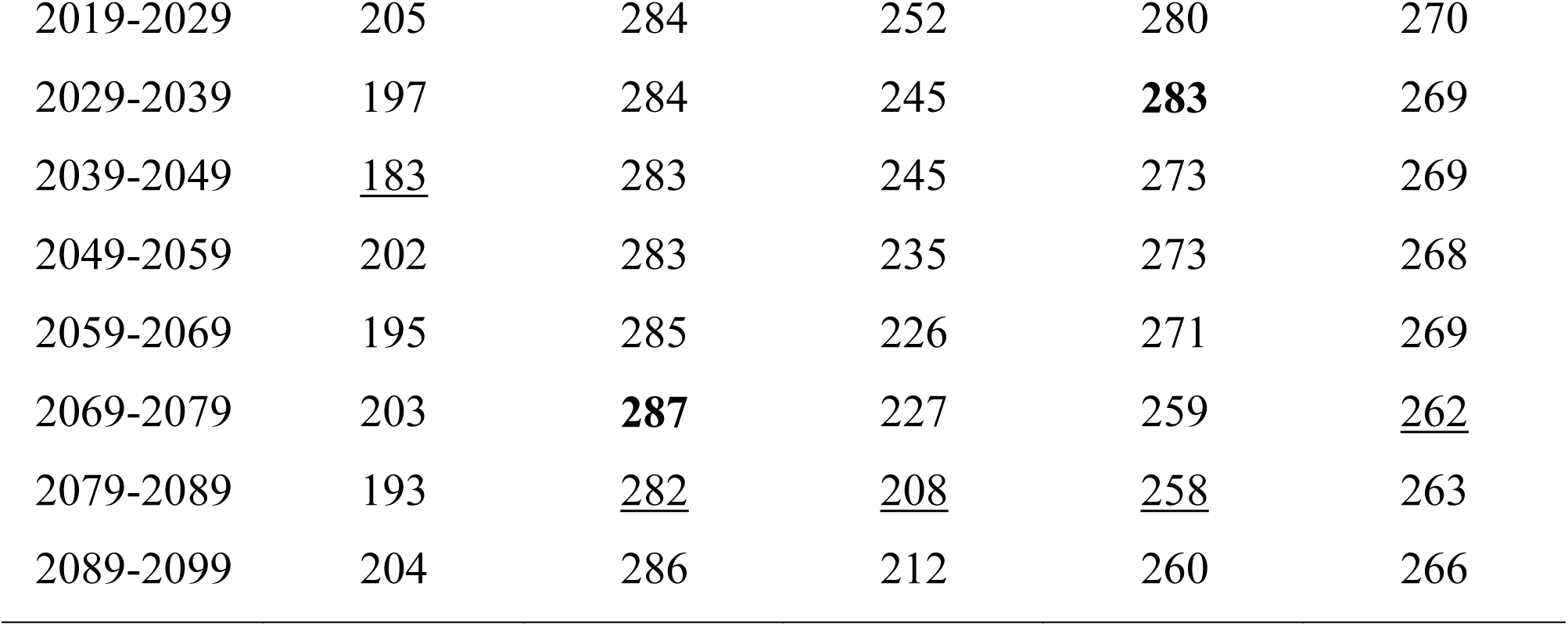
Summary of the changes in the sink/source DoY (Day of Year), considering both climate change scenarios (RCP 2.6 and 6.0) for 5 case studies selected (i.e., Collelongo - IT-Col, Le Bray - FR-LBr, Hyytiälä - FI-Hyy, Bílý Kříž - CZ-Bk1, and Sorø - DK-Sor). In bold values where changes were the highest between the decades while underlined the lowest ones within each forest stand. Data missing for some intervals are because of filtering and data removals to avoid pulsing artefacts e.g. ‘Birch effect’ (see Material and Methods).

**Figure A1.**
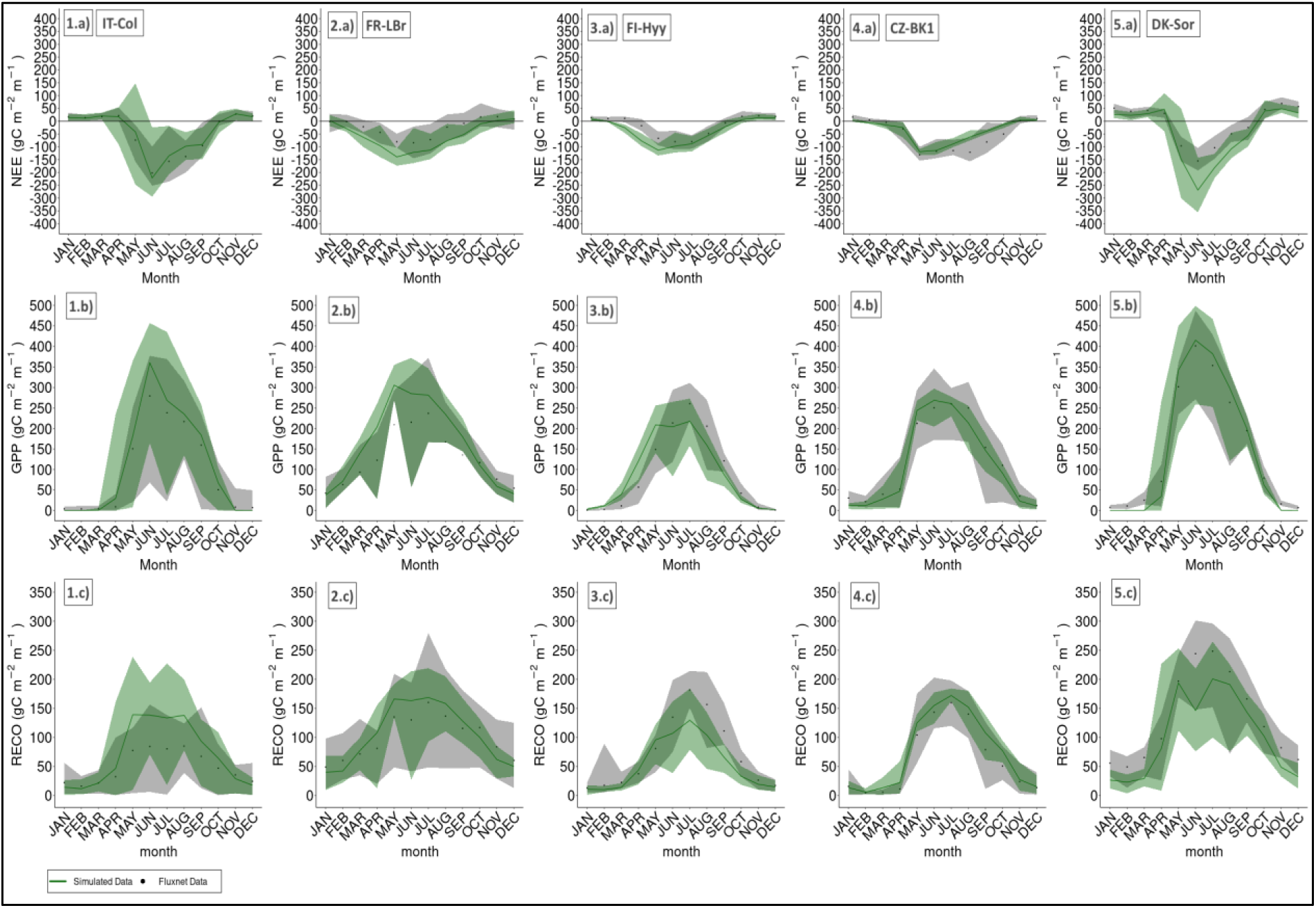
The green lines represent the modeled (a) NEE, (b) GPP and (c) R_eco_ amounts (gC m^—2^ month^—1^) per month for the five selected case studies (i.e., 1) Collelongo – IT-Col, 2) Le Bray - FR-Lbr, 3) Hyytiälä - FI-Hyy, 4) Bílý Křìž - CZ-Bk1, and 5) Sorø - DK-Sor) compared to relative observed data (depicted as black dots) from the Fluxnet2015 Dataset (Pastorello et al., 2020). The lower and upper lines of the shaded area represent, respectively, the minimum and maximum values of the observed and modelled datasets considered.

**Figure A2.**
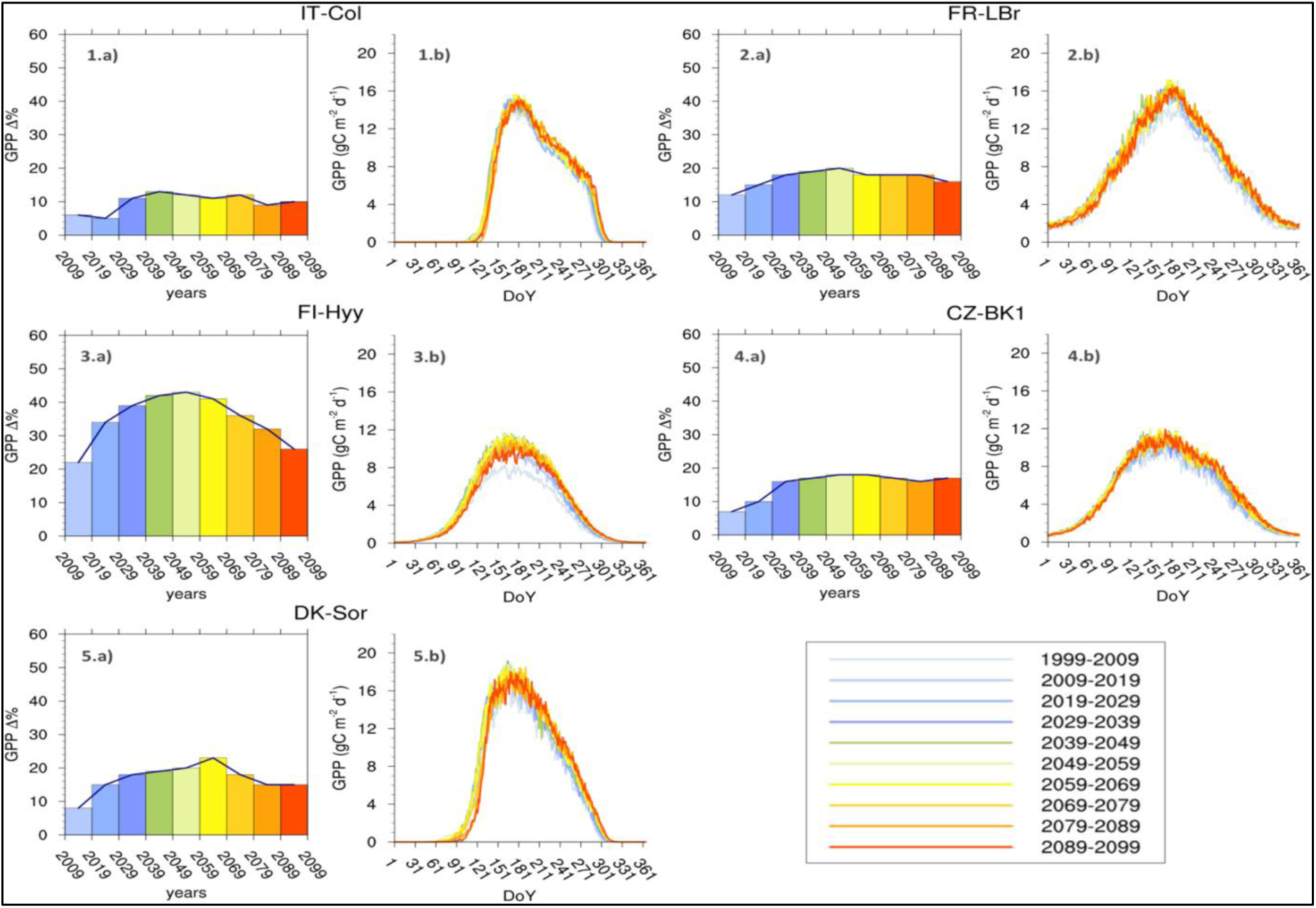
10-year average GPP seasonal cycle under the RCP 2.6 climate scenario for 5 case studies selected, i.e.: 1) Collelongo - IT-Col, 2) Le Bray - FR-LBr, 3) Hyytiälä - FI-Hyy, 4) Bílý Kříž - CZ-Bk1, and 5) Sorø - DK-Sor). The histograms (a) represent the annual GPP variation (%) from the first decade taken as a benchmark of simulation (1999-2009). The xy plots (b) show the Mean Seasonal GPP Cycle of monthly values (gC m^—2^ day^—1^).

**Figure A3.**
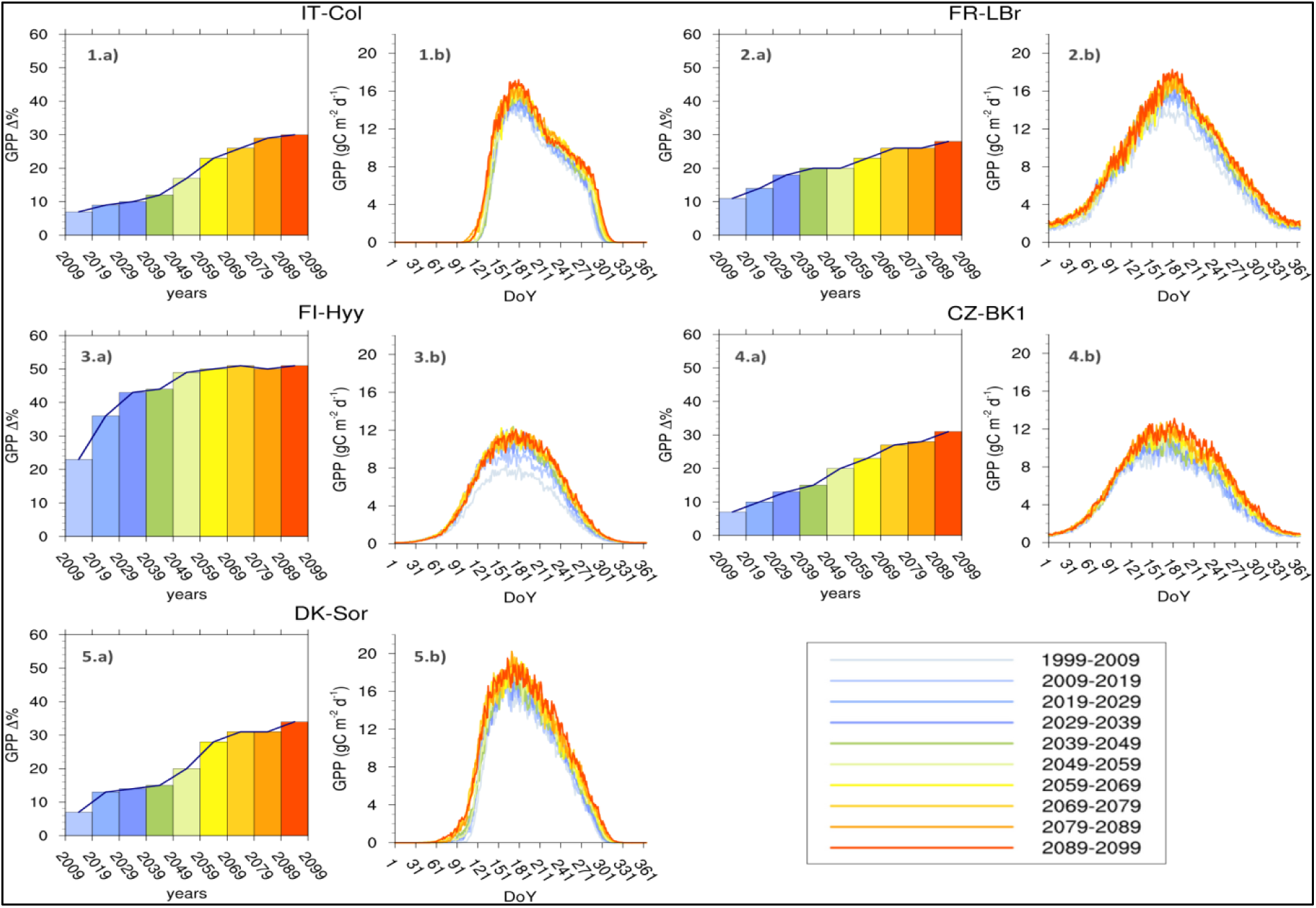
10-year average GPP seasonal cycle under the RCP 6.0 climate scenario for 5 case studies selected, i.e.: 1) Collelongo - IT-Col, 2) Le Bray - FR-LBr, 3) Hyytiälä - FI-Hyy, 4) Bílý Kříž - CZ-Bk1, and 5) Sorø - DK-Sor). The histograms (a) represent the annual GPP variation (%) from the first decade taken as a benchmark of simulation (1999-2009). The xy plots (b) show the Mean Seasonal GPP Cycle of monthly values (gC m^—2^ day^—1^).

**Figure A4.**
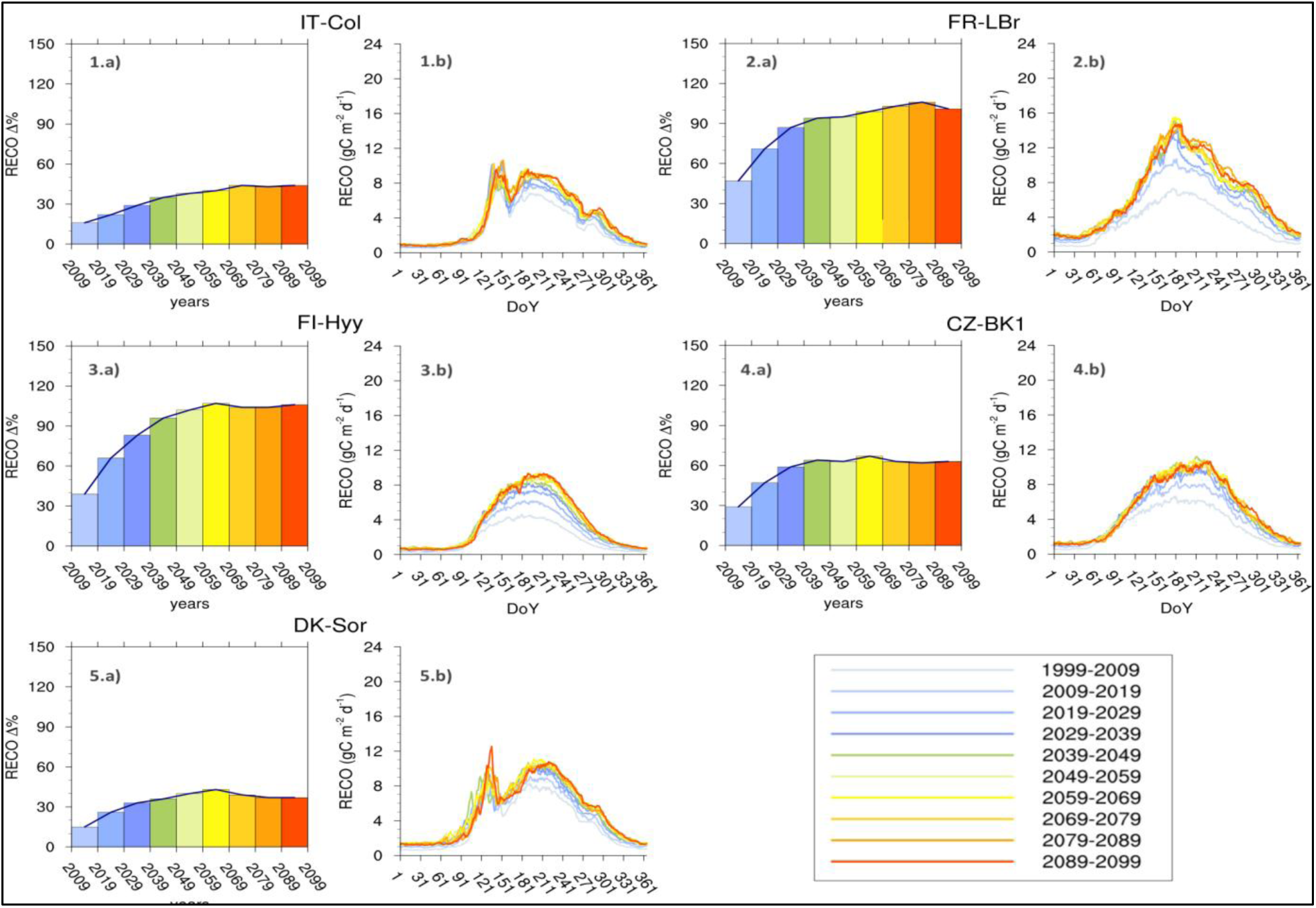
10-year average R_eco_ seasonal cycle under the RCP 2.6 climate scenario for 5 case studies selected, i.e.: 1) Collelongo - IT-Col, 2) Le Bray - FR-LBr, 3) Hyytiälä - FI-Hyy, 4) Bílý Kříž - CZ-Bk1, and 5) Sorø - DK-Sor). The histograms (a) represent the annual R_eco_ variation (%) from the first decade taken as a benchmark of simulation (1999-2009). The xy plots (b) show the Mean Seasonal Reco Cycle of monthly values (gC m^—2^ day^—1^).

**Figure A5.**
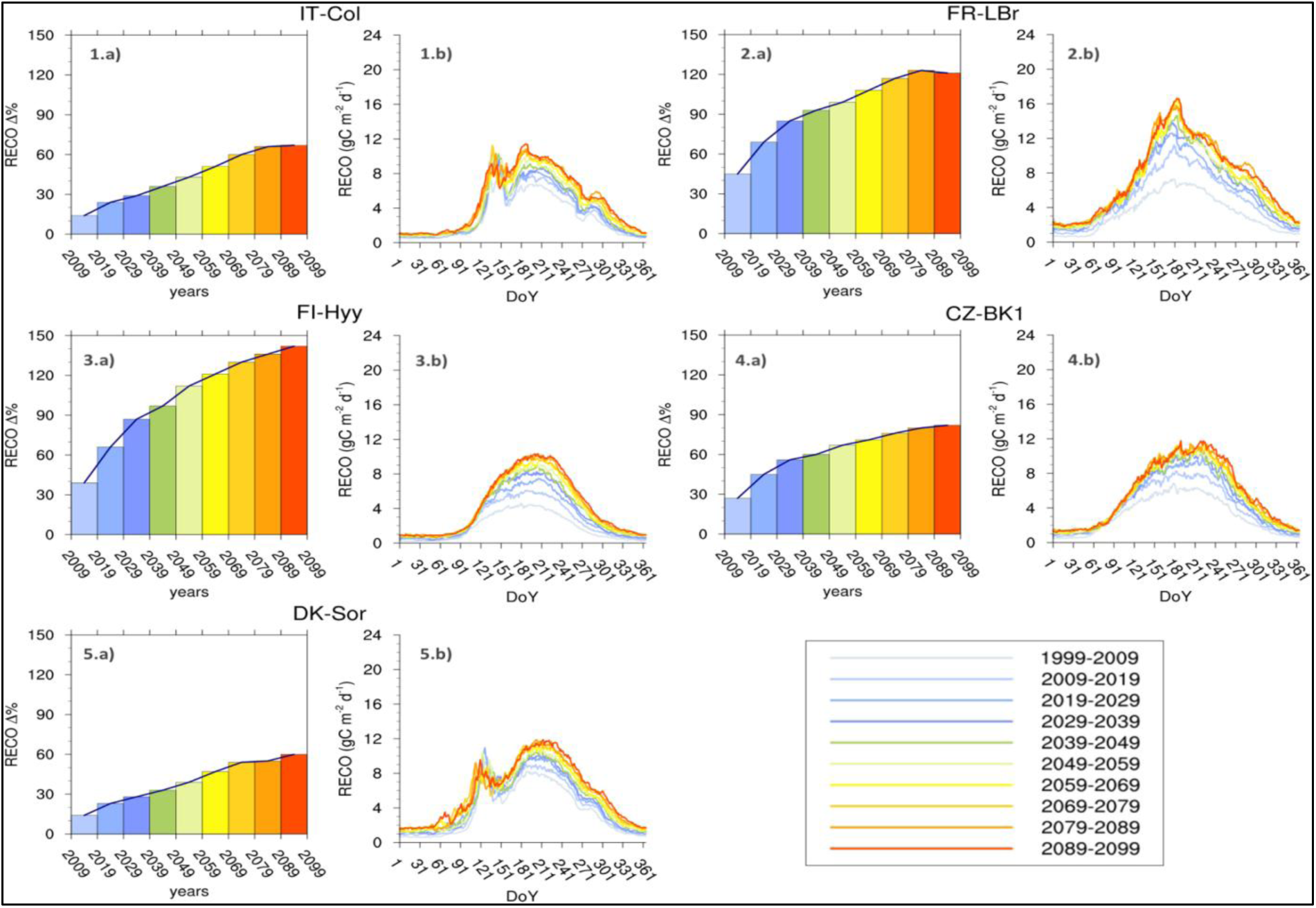
10-year average R_eco_ seasonal cycle under the RCP 6.0 climate scenario for 5 case studies selected, i.e.: 1) Collelongo - IT-Col, 2) Le Bray - FR-LBr, 3) Hyytiälä - FI-Hyy, 4) Bílý Kříž - CZ-Bk1, and 5) Sorø - DK-Sor). The histograms (a) represent the annual R_eco_ variation (%) from the first decade taken as a benchmark of simulation (1999-2009). The xy plots (b) show the Mean Seasonal Reco Cycle of monthly values (gC m^—2^ day^—1^).

